# Statistical analysis and optimality of neural systems

**DOI:** 10.1101/848374

**Authors:** Wiktor Młynarski, Michal Hledík, Thomas R. Sokolowski, Gašper Tkačik

## Abstract

Normative theories and statistical inference provide complementary approaches for the study of biological systems. A normative theory postulates that organisms have adapted to efficiently solve essential tasks, and proceeds to mathematically work out testable consequences of such optimality; parameters that maximize the hypothesized organismal function can be derived *ab initio*, without reference to experimental data. In contrast, statistical inference focuses on efficient utilization of data to learn model parameters, without reference to any *a priori* notion of biological function, utility, or fitness. Traditionally, these two approaches were developed independently and applied separately. Here we unify them in a coherent Bayesian framework that embeds a normative theory into a family of maximum-entropy “optimization priors.” This family defines a smooth interpolation between a data-rich inference regime (characteristic of “bottom-up” statistical models), and a data-limited *ab inito* prediction regime (characteristic of “top-down” normative theory). We demonstrate the applicability of our framework using data from the visual cortex, the retina, and *C. elegans*, and argue that the flexibility it affords is essential to address a number of fundamental challenges relating to inference and prediction in complex, high-dimensional biological problems.

---

Ideas about optimization are at the core of how we approach biological complexity (1–3). Quantitative predictions about biological systems have been successfully derived from first principles in the context of efficient coding (4, 5), metabolic (6, 7), reaction (8, 9), and transport (10) networks, evolution (11), reinforcement learning (12), and decision making (13, 14), by postulating that a system has evolved to optimize some utility function under biophysical constraints. Normative theories generate such predictions about living systems *ab initio*, with no (or minimal) appeal to experimental data. Yet as such theories become increasingly high-dimensional and optimal solutions stop being unique, it gets progressively hard to judge whether theoretical predictions are consistent with data (15, 16), or to define rigorously what that even means (17–19). Alternatively, data may be “close to” but not “at” optimality, and different instances of the system may show variation “around” optima (20, 21), but we lack a formal framework to deal with such scenarios. Lastly, normative theories typically make non-trivial predictions only under quantitative constraints which, ultimately, must have an empirical origin, blurring the idealized distinction between a data-free normative prediction and a data-driven statistical inference.

In contrast to normative theories which derive system parameters *ab initio*, the fundamental task of statistical inference is to reliably estimate model parameters from experimental observations. Here, too, biology has presented us with new challenges. While data is becoming increasingly high-dimensional, it is not correspondingly more plentiful; the resulting curse of dimensionality that statistical models face is controlled neither by intrinsic symmetries nor by the simplicity of disorder, as in statistical physics. To combat these issues and simultaneously deal with the noise and variability inherent to the experimental process, modern statistical methods often rely on prior assumptions about system parameters. These priors either act as statistical regularizers to prevent overfitting or to capture low-level regularities such as smoothness, sparseness or locality (22). Typically, however, their statistical structure is simple and does not reflect the prior knowledge about system function.

Normative theories and inference share a fundamental similarity: they both make statements about parameters of biological systems. While these statements have traditionally been made in opposing “data regimes” (Fig. 1), we observe that the two approaches are not exclusive and could in fact be combined with mutual benefit. To this end, we develop a Bayesian statistical framework that combines data likelihood with an “optimization prior” derived from a normative theory; contrary to simple, typically applied priors, optimization priors can induce a complex statistical structure on the space of parameters. This construction allows us to rigorously formulate and answer the following key questions: (1) Can one derive a statistical hypothesis test for the consistency of data with a proposed normative theory? (2) Can one define how close data is to the proposed optimal solution? *β*) How can data be used to set the constraints in, and resolve the degeneracies of, a normative theory? (4) To what extent do optimization priors aid inference in high-dimensional statistical models? We illustrate the application of these questions and the related concepts to simple model systems, and demonstrate their relevance to real-world data analysis on three diverse examples.

**Fig. 1.**
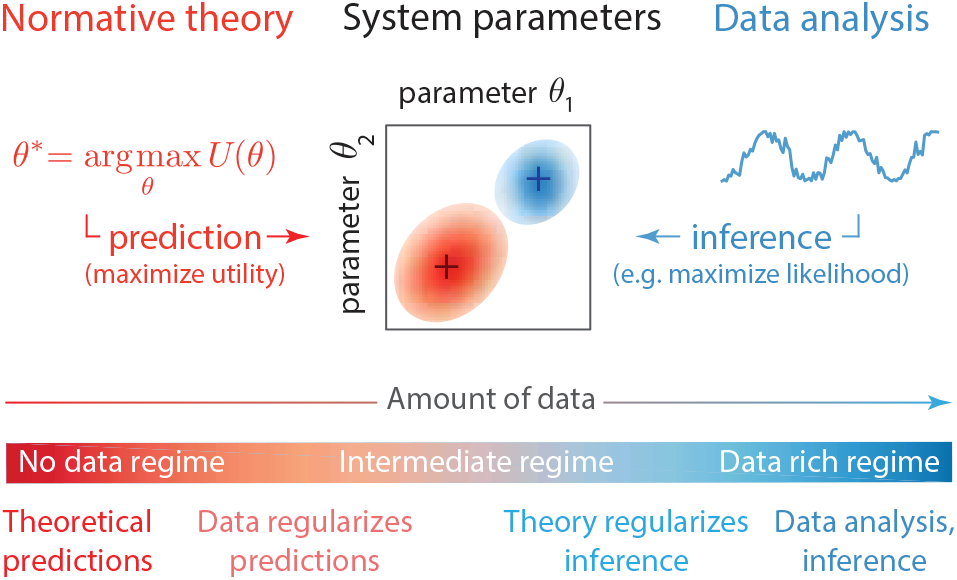
Normative theories and statistical inference. Both approaches make statements about values of system parameters (middle row; center panel). Normative theories predict which parameters would be of highest utility to the system (middle row in red; left panel) without reference to experimental data. Data analysis infers parameter values from experimental observations (middle row in blue; right panel). Large amounts of data support reliable inference of parameters. We consider a continuum of regimes that are applicable with different amounts of data (bottom row).

## Results

### Bayesian inference and optimization priors

Given a probabilistic model for a system of interest, *P* (*x*|*θ*), with parameters *θ*, and a set of *T* observations (or data) 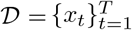, Bayesian inference consists of formulating a (log) posterior over parameters given the data:

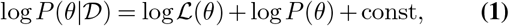

where the constant term is independent of the parameters, 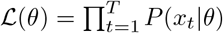 is the likelihood assuming independent and identically distributed observations *x*_*t*_, and *P* (*θ*) is the prior, or the postulated distribution over the parameters in absence of any observation. Much work has focused on how the prior should be chosen to permit optimal inference, ranging from uninformative priors (23), priors that regularize the inference and thus help models generalize to unseen data (24, 25), or priors that can coarse-grain the model depending on the amount of data samples, *T* (26).

Our key intuition will lead us to a new class of priors fundamentally different from those considered previously. A normative theory for a system of interest with parameters *θ* can typically be formalized through a notion of a (upper-bounded) utility function, *U* (*θ*; *ξ*), where *ξ* are optional parameters which specify the properties of the utility function itself. Optimality then amounts to the assumption that the real system operates at a point in parameter space, *θ*^∗^, that maximizes utility, *θ*^∗^(*ξ*) = argmax_*θ*_ *U* (*θ*; *ξ*). Viewed in the Bayesian framework, the assertion that the system is optimal thus represents an infinitely strong prior where the parameters are concentrated at *θ*^∗^, i.e., *P* (*θ ξ*)= *“*(*θ θ*^∗^(*ξ*)). In this extreme case, no data is needed to determine system parameters: the prior fixes their values and typically no finite amount of data will suffice for the likelihood in Eq (1) to move the posterior away from *θ*^∗^. This concentrated prior can, however, be interpreted as a limiting case of a softer prior that “prefers” solutions close to the optimum.

Consistent with the maximum entropy principle put forward by Jaynes (27), we therefore consider for our priors distributions that are as random and unstructured as possible while attaining a prescribed average utility:

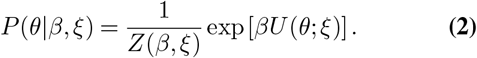

This is in fact a family of priors, whose strength is parametrized by *β*: when *β* = 0, parameters are distributed uniformly over their domain without any structure and in absence of any optimization; as *β* → ∞, parameter probability localizes at the point *θ*^∗^(*ξ*) that maximizes the utility to *U*_max_(*ξ*) (if such a point is unique) irrespective of whether data supports this or not. At finite *β*, however, the prior is “smeared” around *θ*^∗^(*ξ*) so that the average utility, *Ū* (*β, ξ*)= *dθ P* (*θ β, ξ*)*U* (*θ, ξ*) *< U*_max_(*ξ*) increases monotonically with *β*. For this reason, we refer to *β* as the “optimization parameter,” and to the family of priors in Eq (2) as “optimization priors.”

The intermediate regime, 0 *< β <* ∞, in the prior entering Eq (1) is interesting from an inference standpoint. It represents the belief that the system may be “close to” optimal with respect to the utility *U* (*θ*; *ξ*) but this belief is not absolute and can be outweighed by the data: the log likelihood, log ℒ, grows linearly with the number of observations, *T*, matching the roughly linear growth of the log prior with *β*. Varying *β* thus literally corresponds to the interpolation between an infinitely strong optimization prior and pure theoretical prediction in the “no data regime” and the uniform prior and pure statistical inference in the “data rich regime”, as schematized in Fig. 1.

Additional parameters of the utility function, *ξ*, determine its shape in the domain of parameters *θ*. Parameters *ξ* can be known and fixed for a specific theory or, if unknown *a priori*, inferred from the data in a Bayesian fashion. When there are no utility parameters *ξ* to consider, we will suppress them for notational simplicity.

In the following, we apply this framework to a toy model system, a single linear-nonlinear neuron, which is closely related to logistic regression. This example is simple, well-understood across multiple fields, and low-dimensional so that all mathematical quantities can be constructed explicitly; the framework itself is, however, completely general. We then apply our framework to a more complex neuron model and to three experimental data sets. Taken together, these examples demonstrate how the ability to encode the entire shape of the utility measure into the optimization prior opens up a more refined and richer set of optimality-related statistical analyses.

### Example: Efficient coding in a simple model neuron

Let us consider a simple probabilistic model of a spiking neuron (Fig. 2A), a broadly applied paradigm in sensory neuroscience (28–32). The neuron responds to one-dimensional continuous stimuli *x*_*t*_ either by eliciting a spike (*r*_*t*_ = 1), or by remaining silent (*r*_*t*_ = 0). The probability of eliciting a spike in response to a particular stimulus value is determined by the nonlinear saturating stimulus-response function. The shape of this function is determined by two parameters: position *x*_0_ and slope *k* (see Methods).

**Fig. 2.**
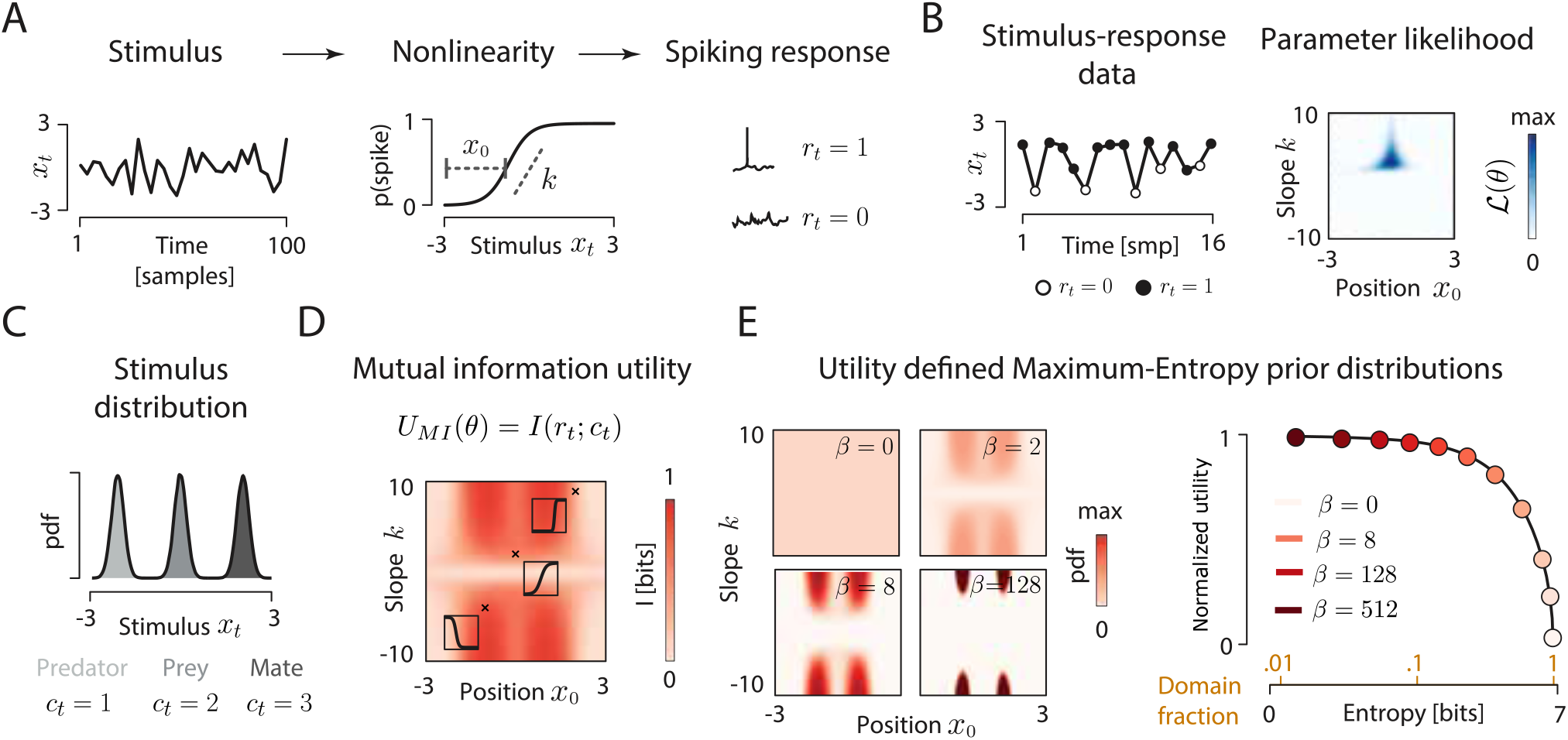
Efficient coding in a toy model neuron and the corresponding optimization prior. **(A)** Model neuron uses a logistic nonlinearity (middle panel) to map continuous stimuli *x*_*t*_ (left panel) to a discrete spiking response *r*_*t*_ (right panel). The shape of the nonlinearity is described by two parameters: slope *k* and offset *x*_0_. **(B)** An example dataset (left panel) consisting of stimulus values (black line) and associated spiking responses (empty circles – no spike, full circles – spike). Likelihood function of the nonlinearity parameters defined by the observed data. Dark blue corresponds to most likely parameter values. **(C)** Distribution of natural stimuli to which the neuron might be adapted. In this example, each mode corresponds to a behaviorally relevant state of the environment: presence of a predator, a prey or a mate. **(D)** Efficient coding utility function, here, the mutual information between neural response *r*_*t*_ and the state of the environment, *c*_*t*_, with stimuli drawn from the distribution in panel C. The amount of information conveyed by the neuron depends on the position and slope of the nonlinearity. Insets depict example nonlinearities corresponding to parameter values marked with black crosses. **(E)** Four maximum-entropy optimization priors over parameters for the neural nonlinearity (left panel). Distributions are specified by the utility of each slope-offset combination. Increasing parameter *β* constrains the distribution (lowers its entropy) and increases the expected utility of the parameters (right panel). Here we plot the normalized utility *U* (*θ*) - see main text for explanation. Orange numbers on the horizontal axis specify the fraction of the entire domain effectively occupied by parameters at given *β*.

Parameters *θ* = {*x*_0_,*k*} fully determine the function of the neuron, yet remain unknown to the external observer. Statistical inference extracts parameter estimates 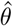 using experimental data 𝒟 consisting of stimulus-response pairs (Fig. 2B, left panel), by first summarizing the data with the likelihood, ℒ (*θ*) (Fig. 2B, right panel), followed either by maximization of the likelihood, 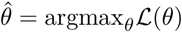 in the maximum-likelihood (ML) paradigm, or by deriving 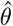 from the posterior, Eq (1), in the Bayesian paradigm.

To apply our reasoning, we must propose a normative theory for neural function, form the optimization prior, and combine it with the likelihood in Fig. 2B, as prescribed by the Bayes rule in Eq (1). An influential theory in neuroscience called “efficient coding” postulates that sensory neurons maximize the amount of information about natural stimuli they encode into spikes given biophysical constraints (5, 31, 33–36). This information-theoretic optimization principle *β*7) has correctly predicted neural parameters such as receptive field (RF) shapes *β*4, 38) and the distribution of tuning curves (17, 39), as well as other quantitative properties of sensory systems (4, 40–44), *ab initio*, from the distribution of ecologically relevant stimuli (2, 34).

To apply efficient coding, we need to specify a distribution from which the stimuli *x*_*t*_ are drawn. In reality, neurons would respond to complex and high-dimensional features of sensory inputs, such as a particular combination of odorants, timbre of a sound or a visual texture, in order to help the animal discriminate between environmental states of very different behavioral relevance (e.g. a presence of a predator, a prey or a mate). To capture this intuition in our simplified setup, we imagine that the stimuli *x*_*t*_ are drawn from a multi-modal distribution, which is a mixture of three different environmental states, labeled by *c*_*t*_ (Fig. 2C). Efficient coding then postulates that the neuron maximizes the mutual information, *I*(*r*_*t*_; *c*_*t*_), between the environmental states, *c*_*t*_, that gave rise to the corresponding stimuli, *x*_*t*_, and the neural responses, *r*_*t*_.

Mutual information, which can be evaluated for any choice of parameters *k, x*_0_, provides the utility function, *U*_MI_(*k, x*_0_)= *I*(*r*_*t*_; *c*_*t*_), relevant to our case; in this simple example, the utility function has no extra parameters *ξ*. Figure 2D shows that *U*_MI_ is bounded between 0 and 1 bit (since the neuron is binary), but does not have a unique maximum. Instead, there are four combinations of parameters that define four degenerate maxima, corresponding to the neuron’s nonlinearity being as steep as possible (high positive or negative *k*) and located in any of the two “valleys” in the stimulus distribution (red peaks in Fig. 2D). Moreover, the utility function forms broad ridges on the parameter surface, and small deviations from optimal points result only in weak decreases of utility. Consequently, formulating clear and unambiguous theoretical predictions is difficult, an issue that has been recurring in the analysis of real biological systems (45, 46).

Given the utility function, the construction of the maximum-entropy optimization prior according to Eq (2) is straightforward. Explicit examples for different values of *β* are shown in Fig. 2E (left panel). Generally, the average utility of the prior monotonically increases as the prior becomes more localized around the optimal solutions, as measured by the decrease in entropy of the prior (Fig. 2E, right panel). This can be interpreted as restricting the system into a smaller part of the parameter domain. If an increase in average utility requires a reduction in entropy by 1 bit, this means that the parameters will be sampled from at most half the available domain.

Before proceeding, we note that our approach depends on several non-trivial choices. First, the fact that system parameterization and the size of the parameter domain can affect Bayesian inferences is well recognized (47) and we discuss how it relates to our case in Supplemental Information (Fig. S1, S2). Second, *β* and the utility function enter the optimization prior of Eq (2) as a product, leaving the scale of each quantity arbitrary. For interpretation purposes we therefore define the normalized utility, *Ũ* = (*Ū* (*β*) − *Ū* (*β* = 0))*/*(*U*_max_ − *Ū* (*β* = 0)), which takes on values between 0 and 1 for non-negative *β*, and is insensitive to linear scaling. We discuss the issue of *β* scaling in Supplemental Information. Third, data and optimality theories could be combined in multiple ways. However combining them via maxent optimization priors enjoys favorable theoretical guarantees that alternative approaches may lack, which we demonstrate in Supplemental Information (Fig. S3, S4). These considerations complete our setup and allow us to address the four questions posed in the Introduction.

### Question 1: Statistical test for the optimality hypothesis

Given a candidate normative theory and experimental data for a system of interest, a natural question arises: Does the data support the postulated optimality? This question is non-trivial for two reasons. First, optimality theories typically do not specify a sharp boundary between optimal and non-optimal parameters, but rather a smooth utility function *U* (*θ*) (Fig. 3A): How should the test for optimality be defined in this case? Second, a finite dataset 𝒟 might be insufficient to infer a precise estimate of the parameters *θ*, but will instead yield a (possibly broad) likelihood surface (Fig. 3B): How should the test for optimality be formulated in the presence of such uncertainty?

**Fig. 3.**
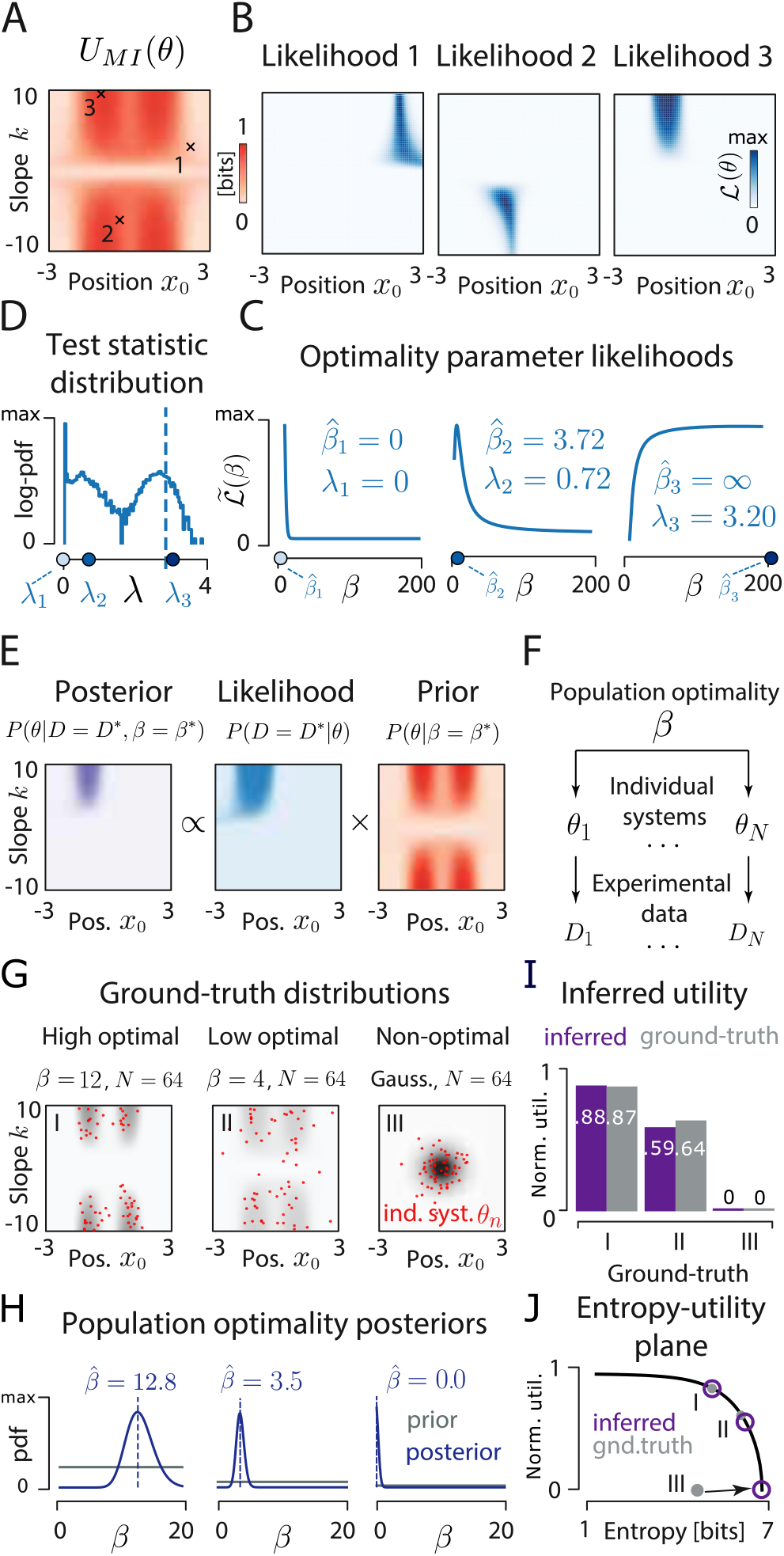
Statistical test (A-D) and inference of the degree (E-J) of optimality. **(A)** Utility function *U*_MI_(*k, x*_0_). Crosses and numbers show the locations of ground truth parameters. **(B)** Likelihood of the nonlinearity parameters obtained from 20 stimulus–response (*x*_*i*_, *r*_*i*_) pairs. The three examples correspond to three ground truth parameter values (black crosses in A), and are ordered by increasing utility. **(C)** Marginal likelihood of the optimality parameter *β*, 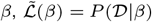, corresponding to data in A. Maximum likelihood estimates 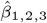 (blue circles) indicate that the data would be most probable with no preference for high utility *U*_MI_ (left panel, 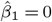 – note that we do not allow negative 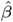), some preference for high *U*_MI_ (middle panel, 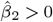 finite) and strong preference for high *U*_MI_ (right panel, 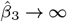 blue circle displayed at *β* = 200 for illustration purposes). Likelihood ratio statistic *λ*_1,2,3_ compares the marginal likelihood of *β* at *β* = 0 vs 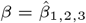 (see Methods). **(D)** Null distribution of the test statistic *λ*. Point mass at *λ* = 0 corresponds to cases where the maximum likelihood optimality parameter is zero, 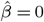. High values of *λ* are evidence against the null hypothesis that *β* = 0, and hence support optimality. Dashed vertical line represents *p* = 0.05 significance threshold, blue circles show *λ*_1,2,3_. Only *λ*_3_ crosses the threshold, indicating significant preference for high utility parameters. **(E)** Posterior over nonlinearity parameters, inferred for a single system with a utility-derived prior at fixed optimality parameter, *β* = *β*^∗^. **(F)** A hierarchical model of a population of optimized systems. Population optimality parameter *β* controls the distribution of parameters for individual systems (*n* = 1,…,*N*), *θ*_*n*_, which give rise to observed data, *D*_*n*_. **(G)** Nonlinearity parameters (64 red dots per distribution) sampled from three different ground truth distributions (denoted by roman numerals in panels G-J): a strongly optimized population (*β* = 12; left), a weakly optimized population (*β* = 4; middle), a non-optimal distribution (Gaussian distribution; right). For each model neuron *θ*_*n*_, data *D*_*n*_ consists of 100 stimulus-response pairs. **(H)** Results of hierarchical inference. Posteriors over *β* (purple lines) and MAP estimates, 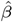 (dashed purple lines) were obtained using simulated data from G. Priors (gray lines) were uniform on the [0, 20] interval. **(I)** Normalized utility *Ũ*. Estimated values (purple bars) closely match ground truth (gray bars). **(J)** Entropy and normalized utility of ground truth distributions (gray, filled circles) and inferred distributions parametrized by 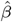 (purple, empty circles).

Here we devise an approach to address both issues. The basis of our test is a null hypothesis that the system is not optimized, i.e., that its parameters have been generated from a uniform random distribution on the biophysically accessible parameter domain. This distribution is exactly the optimization prior *P* (*θ*|*β* = 0). The alternative hypothesis states that the parameters are drawn from a distribution *P* (*θ*|*β*) with *β>* 0. To discriminate between the two hypotheses, we use a likelihood ratio test with the statistic λ, which probes the overlap of high-likelihood and high-utility parameter regions. Specifically, we define the marginal likelihood of *β* given data, 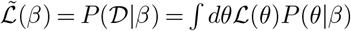 (Fig. 3C), and then define *λ* as the log ratio between the maximal marginal likelihood, max 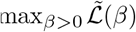, and the marginal likelihood under the null hypothesis, 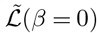 (see Methods).

The test statistic *λ* has a null distribution that can be estimated by sampling (Fig. 3D), with large *λ* implying evidence against the null hypothesis; thus, given a significance threshold, we can declare the system to show significant degree of optimization, or to be consistent with no optimization. This is different from asking if the system is “at” an optimum: such a narrow view seems too restrictive for complex biological systems. Evolution, for example, might not have pushed the system all the way to the biophysical optimum (e.g., due to mutational load or because the adaptation is still ongoing), or the system may be optimal under utility function or resource constraints slightly different than those postulated by our theory (21). Instead, the proposed test asks if the system has relatively high utility, compared to the utility distribution in the full parameter space.

While principled, this hypothesis test is computationally expensive, since it entails an integration over the whole parameter space to compute the marginal likelihoods, 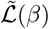, as well as Monte Carlo sampling to generate the null distribution. The first difficulty can be resolved when the number of observations *T* is sufficient such that the likelihood of the data, ℒ (*θ*), is sharply localized in the parameter space; in this case the value of the utility function at the peak of the likelihood itself becomes the test statistic and the costly integration can be avoided (see Methods). The second difficulty can be resolved when we can observe many systems and collectively test them for optimality; in this case the distribution of the test statistic approaches the standard *χ*^2^ distribution (see Methods).

### Question 2: Inferring the degree of optimality

Hypothesis testing provides a way to resolve the question whether the data provides evidence for system optimization or not (or to quantify this evidence with a p-value). However, statistical significance does not necessarily imply biological significance: with sufficient data, rigorous hypothesis testing can support the optimality hypothesis even if the associated utility increase is too small to be biologically relevant. Therefore, we formulate a more refined question: How strongly is the system optimized with respect to a given utility, *U* (*θ*)? Methodologically, we are asking about the value of the optimization parameter, *β*, that is supported by the data 𝒟. In the standard Bayesian approach, all parameters of the prior are considered fixed before doing the inference; the prior is then combined with likelihood to generate the posterior (Fig. 3E). Our case corresponds to a hierarchical Bayesian scenario, where *β* is itself unknown and of interest. In the previous section we chose it by maximizing the marginal likelihood, 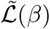, to devise a yes/no hypothesis test. Here, we consider a fully Bayesian treatment, which is particularly applicable when we observe many instances of the same system. In this case, we interpret different instances (e.g., multiple recorded neurons) as samples from a distribution determined by a single population optimality parameter *β* (Fig. 3F) that is to be estimated. Stimulus-response data from multiple neurons are then used directly to estimate a posterior over *β* via hierarchical Bayesian inference.

To explore this possibility, we generate parameters *θ*_*n*_ of *n* = 1,…,*N* model neurons from three different distributions: strongly optimized (*β* = 12; Fig. 3G, left panel), weakly optimized (*β* = 4; Fig. 3G, middle panel) and non-optimal (Gaussian distribution of parameters; Fig. 3G, right panel). For each of the three examples, we simulate stimulus-response data for all neurons and use these data in a standard hierarchical Bayesian inference to compute posterior distributions over the population optimality parameter, *β* (Fig. 3H; see Methods).

Following hierarchical inference, we can interpret the inferred population optimality parameter 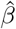, by mapping it onto normalized utility (cf. Fig. 2E). This reports optimality on a [0, 1] scale, with 1 corresponding to the maximum achievable utility *U*_max_ and thus a fully optimal system, and 0 corresponding to the average utility under random parameter sampling, *Ū* (*β* = 0). Normalized utility for the three examples is shown in Fig. 3I.

Our framework enables us to draw inferences about optimality which are not possible otherwise. For example, in addition to estimating the normalized utility, we can also quantify how restrictive the optimization needs to be in order to achieve that level of utility. This restriction is measured by the entropy associated with 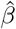 (Fig. 3J). In example I from Fig. 3G-I, 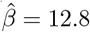 is associated with a decrease in entropy of about 1.75 bits compared to *β* = 0, meaning that nonlinearity parameters are effectively restricted to a fraction about 2^*−*1.75^ ≈ 0.3 of the parameter domain. Example III with 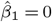 is consistent with a high-entropy optimization prior and indicates almost no parameter space restriction. This is despite the fact that the actual parameters were sampled from a Gaussian highly concentrated (i.e., with low entropy) in the parameter space *β* but not in a region of high utility. This mismatch suggests that such a system could be optimized for a different utility function or shaped by other constraints. The system could also be anti-optimized, i.e. prefer negative values of *Ũ*, which could easily be identified by permitting negative *β* values during inference. Another clear benefit of the probabilistic framework is the possibility of computing uncertainty estimates of *β* and the associated utility and entropy.

### Question 3: Data resolves ambiguous theoretical predictions

Predictions derived from optimality theories can be non-unique and ambiguous. This ambiguity can manifest itself in different ways.

The *first kind of ambiguity* results from the existence of multiple maxima of the utility function. Before formulating statistical questions, it is important to pause and clarify the underlying biological context: Could different observed instances of the system freely sample from all utility maxima (as in Fig 3G, example I), or is a single maximum relevant, perhaps because it is the only one that nature realized by evolutionary adaptation? In the later case, the first task of statistical analysis is to identify that single maximum. For low-dimensional systems, this ambiguity can be resolved trivially: in our toy model, for example, a few data points suffice to zero in on one of the four degenerate utility maxima (Fig. S5). In contrast, in high-dimensional parameter spaces the task of finding the “closest optimum” is non-trivial (15) and could be aided by sampling methods derived from optimization priors, which is a topic for further research.

The *second kind of ambiguity* results from system parameters which enter the utility function, but are unconstrained by the optimization theory in question. Such parameters limit the performance of the whole system, with the utility typically achieving its global maximum when they take on extremal values (e.g., *±* ∞, 0, etc.); yet, these extremal values often correspond to physically implausible scenarios (infinite averaging time or energy consumption, zero noise, instantaneous response time, etc.). Optimization theory cannot make a non-trivial prediction about these parameters, so they must either be fixed *a priori* based on known external constraints, or inferred from data simultaneously with the optimization of the remaining parameters. An additional subtlety comes into play when we analyze multiple instances of a system (e.g., neurons): either each individual neuron has its own value of the constraint parameter, to be determined from data (which we address in the following paragraph), or all neurons share a single value of the constraint that needs to be inferred jointly (statistically equivalent to the third kind of ambiguity in the utility function, which we address subsequently).

In our model, the nonlinearity slope *k* is unconstrained by optimization: mutual information increases monotonically as |*k*| → ∞ (Fig. 4A). This corresponds to vanishing noise in neural spiking. Since such noise cannot physically vanish, we must change the interpretation of the utility function, *U*_MI_(*θ*), and evaluate it only over positions *x*_0_, while treating the slope *k* as a constraint to be fit from data*β*which we indicate by writing *U*_MI_(*x*_0_; *k*). Here, slope *k* determines the entire shape of the utility function (Fig. 4B). Unreliable neurons with a small slope have a unique optimal position *x*_0_ = 0, while for neurons with large *k* the utility is bimodal, with optimal positions separating peaks of the stimulus distribution. As before, we can infer both parameters for a “noisy” (Case I) and “precise” (Case II) simulated neuron (Fig. 4C); this time, however, the optimization prior acts only on *x*_0_, while the prior over slope |*k*| remains uniform. To properly assess optimality, we must normalize the utility by the maximal utility achievable at the estimated value of 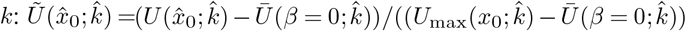. In both cases, the relative utility exceeds 0.9 (Fig. 4C). Because theoretical predictions now depend on the biophysical constraint*β*which itself is a free parameter adjustable separately for each system instance*β*high values of normalized utility can be achieved by neurons with very different *x*_0_.

**Fig. 4.**
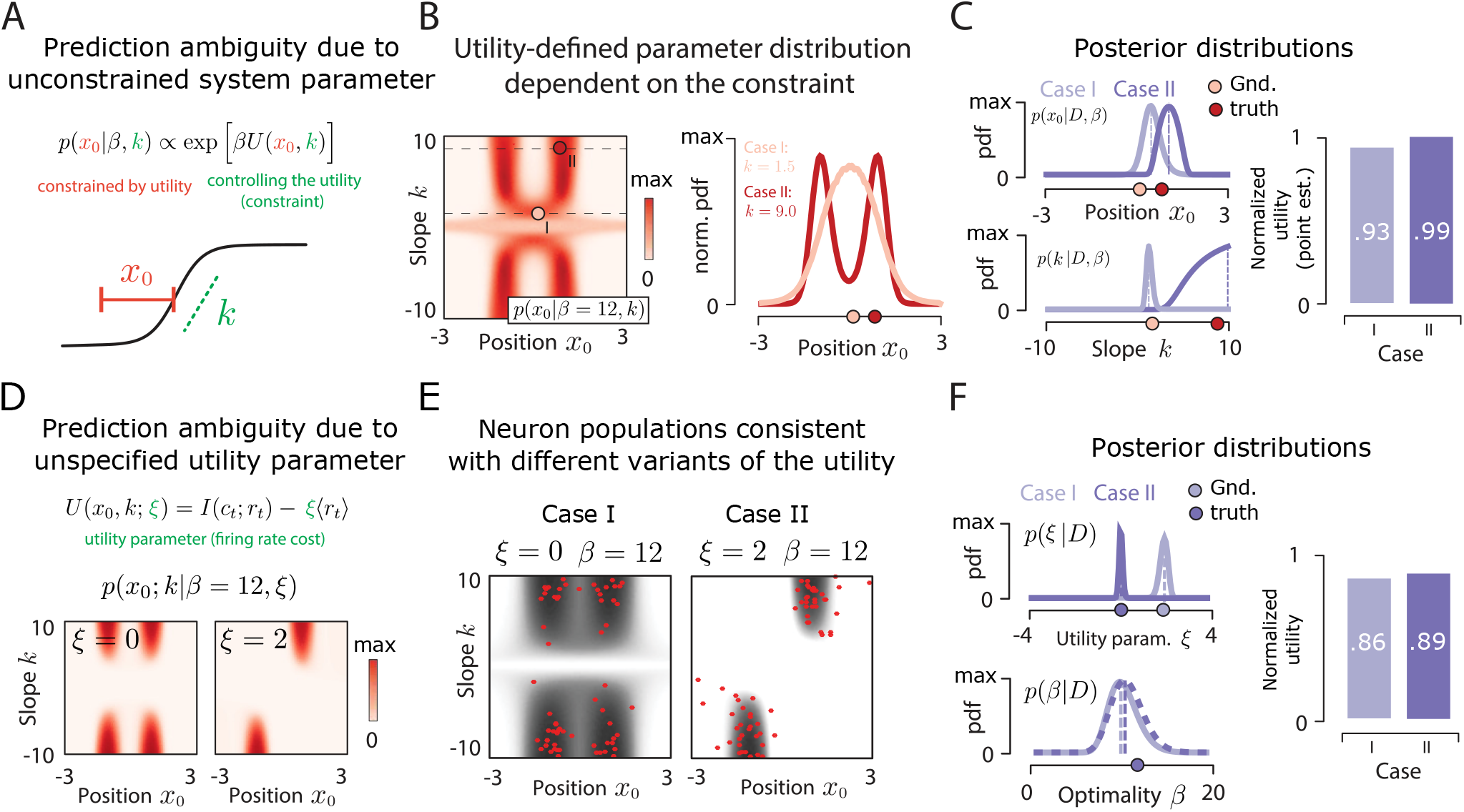
Resolving ambiguities of theoretical predictions. **(A)** Prediction ambiguity due an unconstrained system parameter. Utility is evaluated over the position parameter *x*_0_ (red), with the slope parameter *k* (green) interpreted as an externally imposed biophysical constraint. *k* is inferred from data for each neuron separately; for different *k*, optimality may predict different optimal positions, *x*_0_. **(B)** Optimization priors for *x*_0_ are conditional maxent distributions over *x*_0_ parametrized by values of *k* (rows of the matrix), here at fixed *β* = 12 (left). Distributions over *x*_0_ for two example values of *k* (dashed black lines at left) are displayed in the right panel, with optimal *x*_0_ values marked (pink and red circles for cases I and II, respectively). **(C)** Posteriors over the position (*x*_0_, left column, top) and the slope (*k*, left column, bottom) parameters, estimated for cases I and II (light and dark purple lines, respectively; dashed lines – MAP estimates), by marginalizing the joint posterior. Ground-truth values are marked with circles. Normalized utility of *x*_0_, relative to the maximal utility for *k* inferred separately for cases I and II. **(D)** Prediction ambiguity due to an unspecified utility function. Utility prefers high mutual information *I* at a low average firing rate ⟨*r*⟩, with an unknown trade-off parameter *ξ*. Optimization prior with no firing rate constraint (left, *ξ* = 0) shows four degenerate maxima; the constraint (right, *ξ* = 2) partially lifts the degeneracy. **(E)** Two ground truth distributions (gray) corresponding to different values of the firing rate constraint *ξ*. Red dots denote *N* = 64 sample neurons. **(F)** Posteriors over the firing rate constraint *ξ* (left column, top) and the optimality parameter *β* (left column, bottom), estimated for cases I and II (light and dark purple lines, respectively; dashed lines – MAP estimates), by marginalizing the joint posterior. Ground-truth values are marked with circles. Normalized utilities computed for *ξ* inferred separately for cases I and II.

The *third kind of ambiguity* arises when the utility function itself depends on additional parameters, *ξ*. The mutual information utility *U*_MI_ of our toy model can be extended by considering the cost of neural spiking, resulting in a new compound function, *U* (*x*_0_,*k*; *ξ*)= *U*_MI_(*x*_0_,*k*) *− ξ*⟨*r*_*t*_⟩, with the trade-off parameter *ξ*. Increasing *ξ* changes the shape of the new utility function (Fig. 4D). Given multiple instances of a biological system (Fig. 4E), we can ask about the most likely form of *U* (i.e., the single value of *ξ* shared across all instances of the system), together with the most likely value of the optimization parameter, *β*. This problem is solved by hyperparameter inference, which generates joint posteriors and MAP estimates of *β* and *ξ* (Fig. 4F). Here, too, the normalized utilities are defined relative to the inferred value of *ξ* and can thus be comparable even when the underlying utility functions are substantially different.

### Question 4: Optimization priors improve inference for high-dimensional problems

Here, we extend our toy model neuron with 2 parameters to a more realistic case with hundreds of parameters. We focus on a Linear-Nonlinear-Poisson (LNP) model *β*0), whose responses to natural image stimuli are determined by a linear filter (also referred to as a receptive field - RF) - *ϕ* ∈ ℝ^16×16^ (Fig. 5A). The purpose of this exercise is to show the tractability of our approach and the power of optimization priors for high-dimensional inference problems. Inference of neural filters, *ϕ*, from data is a central data analysis challenge in sensory neuroscience, making our example practically relevant.

**Fig. 5.**
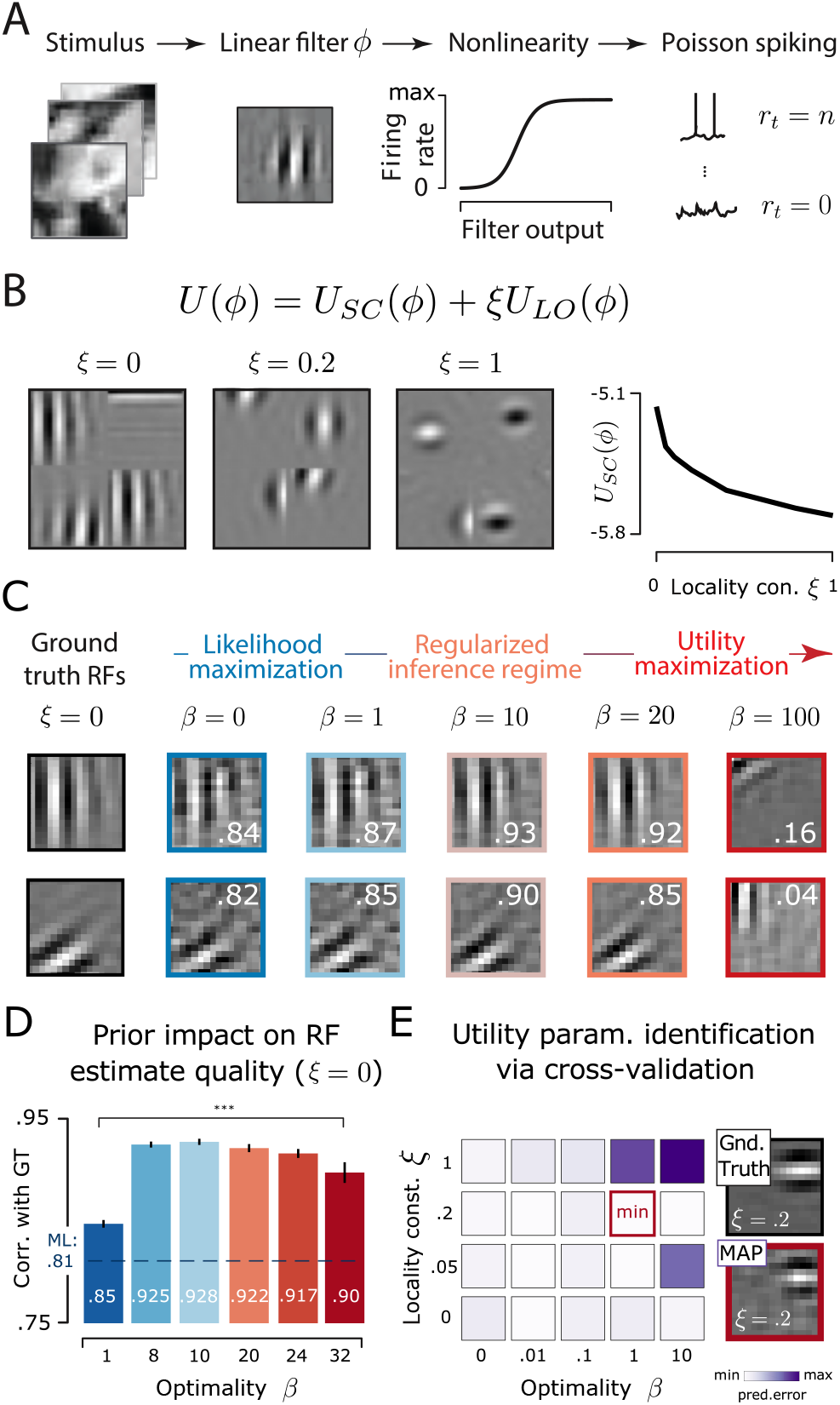
Optimality priors improve inference of high-dimensional receptive fields. **(A)** Linear-nonlinear-Poisson (LNP) neuron responding to 16 × 16 pixel natural image patches, *x*_*t*_. Stimuli are projected onto a linear filter *ϕ*, which transforms them via logistic nonlinearity into average firing rate of Poisson spiking, *r*_*t*_. **(B)** Receptive fields optimized for maximally sparse response to natural stimuli with a locality constraint *ξ*. First three panels on the left display 2 × 2 example filters optimized at increasing *ξ*. Rightmost panel shows the decrease in average sparse utility of filters with increasing *ξ*. **(C)** MAP estimates of two optimally sparse filters (*ξ* = 0) obtained with optimality prior of increasing strength *β*. White digits denote correlation with the corresponding ground truth. **(D)** Average correlations of *N* = 100 filter estimates with the ground truth as a function of prior strength *β* for locality constraint *ξ* = 0. Dashed blue line denotes the average correlation for ML estimates. MAP estimate correlations are significantly higher than ML estimate correlations (t-test; *** denotes *p <* 0.001). Error bars denote standard errors of the mean. **(E)** Identification of prior strength *β* and locality constraint *ξ* via cross-validation. Left panel, cross-validation errors in predicting withheld neural responses for a range of *β* and *ξ* values (heatmap). Parameter combination resulting in minimal error is marked with a red frame. Top right, a ground truth filter optimized with *ξ* = 0.2. Bottom right, MAP estimate of the filter, obtained with correctly identified values for *β* and *ξ*.

Experimentally observed filters *ϕ* in the visual cortex have been suggested to maximize the sparsity of responses *s*_*t*_ to natural stimuli *β*4). A random variable is sparse when most of its mass is concentrated around 0 at fixed variance. These experimental observations have been reflected in the normative model of sparse coding, in which maximization of sparsity has been hypothesized to be beneficial for energy efficiency, flexibility of neural representations, and noise robustness *β*8, 48). Filters optimized for sparse utility *U*_SC_(*ϕ*) (see Methods) are oriented and localized in space and frequency (Fig. 5B, leftmost panel) and famously resemble RFs of simple cells in the primary visual cortex (V1). A significant fraction of neural RFs, however, differ from optimally sparse filters (49), perhaps due to the existence of additional constraints. One possible constraint is spatial locality, which leads to suboptimally sparse filters that increasingly resemble localized blobs (50), as shown in Fig. 5B. In our framework, sparse coding utility *U*_SC_ and locality *U*_LO_ combine into a single utility function with a parameter *ξ* that specifies the strength of the locality constraint. We wondered whether an optimization prior based on sparsity, even in the presence of an additional constraint of unknown strength, could successfully regularize the inference of linear filters, *ϕ*.

We first consider a scenario where the locality constraint is known *a priori* to equal zero. We simulate spike trains of 100 model neurons optimized under sparse utility *U*_SC_ responding to a sequence of 2000 natural image patches (see Methods for details). Using these simulated data we infer the filter estimates, 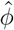, using Spike Triggered Average (STA) (18, 51), which under our assumptions are equivalent to the maximum likelihood (ML) estimates *β*0) (see Methods). STAs computed from limited data recover noisy estimates of neural filters (Fig. 5C; column second from the left).

Can sparse coding provide a powerful prior to aid inference of high-dimensional filters? Using our sparse coding utility, *U*_SC_(*ϕ*), we formulate optimization priors for various values of *β* and compute maximum-a-posteriori (MAP) filter estimates 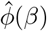 from simulated data (Fig. 5C; four right-most columns; see Methods for details). Increasing values of *β* interpolate between pure data-driven ML estimation (Fig. 5C, second column from the left) that ignores the utility, and pure utility maximization (Fig. 5C, right column) at very high *β* = 10^2^ where the predicted filters become almost completely decoupled from data; these two regimes seem to be separated by a sharp transition. For intermediate *β* = 1, 10, 20, MAP filter estimates show a significant improvement in estimation performance relative to the ML estimate (Fig. 5D).

We next consider a scenario where the locality constraint is not known *a priori*, but can be identified together with the prior strength *β* using cross-validation (52), as described in Question 3. To this end, we simulate responses of a single neuron whose filter was optimized with the locality constraint *ξ* = 0.2 (Fig. 5E, “Gnd. Truth”). We then use a subset of 1800 out of 2000 stimulus-response pairs to compute the MAP estimate of the filter using a range of *β* and *ξ* values. Each MAP estimate of the filter is used to compute the prediction error for neural responses over withheld portion of the data. Cross-validation correctly identifies the true *ξ* and the optimal *β* values which minimize the prediction error (Fig. 5E); the resulting filter estimate (Fig. 5E, “MAP”) closely resembles the ground truth.

Optimization priors achieve a boost in performance because they quantitatively encode many characteristics we ascribe to the observed receptive fields (localization in space and bandwidth, orientation), which the typical regularizing priors (e.g., L2 or L1 regularization of *ϕ* components) will fail to do. While hand-crafted priors designed for receptive field estimation can capture some of these characteristics (18, 53), optimization priors grounded in the relevant normative theory represent the most succinct and complete way of summarizing our prior beliefs. For flexible optimization priors whose strength and additional parameters are set by cross-validation, one might expect that the postulated optimality theory need not be exactly correct to aid inference, so long as it captures *some* of the statistical regularities in the data.

### Application 1: Receptive fields in the visual cortex

Here we analyze receptive fields of neurons in the primary visual cortex (V1) of the Macaque monkey (49) (Fig. 6A). This system is a good test case, for which multiple candidate optimality theories were developed and tested against data (34, 38, 54, 55). As in the example of Fig. 5, we focus on sparse coding using utility *U*_SC_, which prioritizes RFs localized in space and frequency (Fig. 6B; see Methods). An alternative utility prioritizing slow features is presented in Supplemental Information (Fig. S6).

**Fig. 6.**
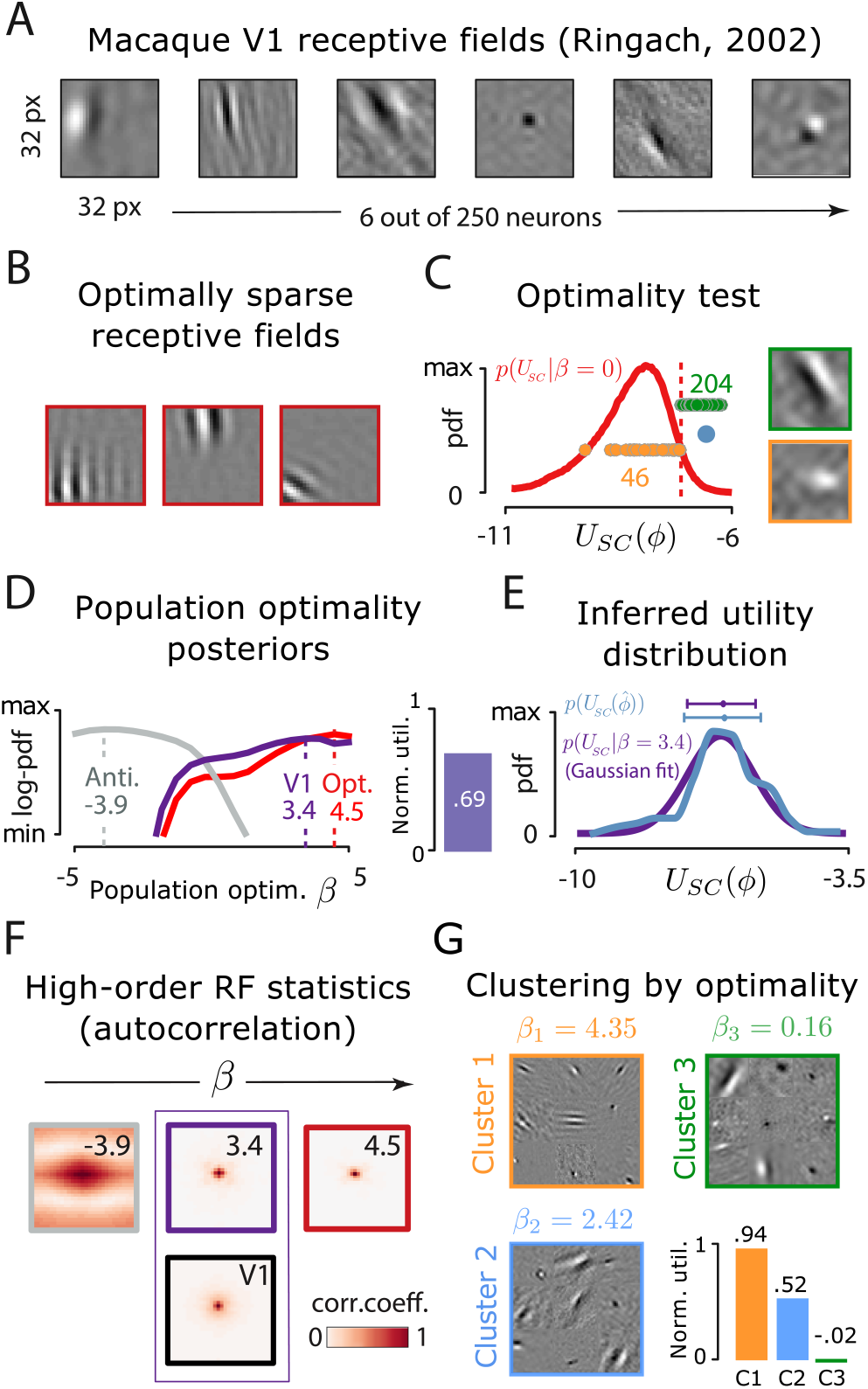
Optimality of V1 receptive fields. **(A)** Six example receptive fields (RFs) from Macaque visual cortex (courtesy of Dario Ringach (49)). **(B)** Example simulated RFs optimized for sparsity. **(C)** Null distribution of utility values used to test for optimality under sparse utility and the 95-percentile significance threshold (red dashed line). Significant (green) and non-significant (orange) receptive fields denoted with dots (x axis is truncated for visualization purposes); example RFs are shown in frames of matching colors. Blue dot shows the average RF utility (99.6^th^ percentile of the null distribution). **(D)** Approximate log-posteriors over population optimality parameter *β* derived from 250 RFs estimates (purple line), 250 maximum-utility filters (red line) and 250 minimal-utility filters (gray line). Dashed lines mark MAP estimates. **(E)** Empirical distribution of RF utilities (blue line) compared with utility distribution consistent with the inferred 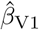 (purple line). **(F)** Spatial autocorrelation of RFs predicted for different *β* values (reported in top-right corner of each panel, cf. inferred values in D). Note a good match between data-derived RF autocorrelation (black frame) and the predicted autocorrelation at the inferred 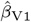 (purple frame). **(G)** Three clusters with different *β*, learned with a MaxEnt mixture-model. For each cluster, 3 × 3 sample receptive fields are displayed, together with the corresponding normalized utility values in the bottom-right panel.

We first ask whether RFs of individual neurons support the optimality hypothesis, as in Question 1. Given the high-quality of RFs estimates, costly marginalization of the likelihood can be avoided and the utility of estimated RFs can be used directly as a test statistic. To construct the null distribution for the test, we sample 10^6^ random filters consistent with optimization prior *P* (*ϕ*|*β* = 0), and declare the 95th percentile to be the optimality threshold (Fig. 6C). As expected, a large majority (204 neurons, green dots / example frame in Fig. 6C) of V1 neurons pass the optimality threshold, with 46 neurons failing the test (orange dots / example frame in Fig. 6C).

We next ask whether all RFs can be used together to quantify the degree of population optimality, as in Question 2. We estimate approximate posteriors over parameter *β* via rejection sampling (see Methods), using all RFs in the population (Fig. 6D, purple line). For comparison, we also compute posteriors using 250 utility-maximizing and 250 utility-minimizing filters (Fig. 6D, red and gray lines, respectively). MAP estimates of *β* obtained with simulated maximal and minimal utility RFs provide a reference for the interpretation of *β* estimated from real data. This estimate, 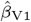, is very close to the parameter value of the optimally sparse filters, implying high degree of optimization. The normalized utility is 0.69, implying a significant, yet not complete, degree of optimization.

Since population optimality *β* parametrizes the entire distribution of receptive fields, inferring *β* allows us to make predictions inaccessible by other means. For example, given the inferred degree of optimality, we predict the entire distribution of utility values (not only its mean) across neurons. In principle, the predicted distribution (or its higher-order moments, e.g., variance) could deviate from the empirically observed distribution, if the real system were adapted to a different utility or set of constraints. For V1 neurons, the predicted and empirical sparse utility distributions are very similar (Fig. 6E).

Another prediction concerns the correlation between system parameters, in our case, RF shapes. Different values of *β* predict very different spatial autocorrelation functions of RFs (Fig. 6F), with the prediction at inferred *β* resembling the data-derived autocorrelation better than the alternative or extremal *β* values. These examples demonstrate that once the single parameter *β* is inferred, the optimality framework makes quantitative, rigorous, and parameter-free predictions of non-trivial statistics that can be directly tested against data. Our framework can also be used to dissect sources of deviation from optimality. We fit a mixture model, where each mixture component was parametrized by a separate value of *β* (Fig. 6G; see Methods). This procedure clusters the RFs into three groups spanning a broad range of utility values. The largest cluster (135 RFs) achieves a nearly maximal normalized utility of 0.94; neurons in this cluster all passed the significance test in Fig. 6C. The existence of second- and third-largest clusters (95 RFs, normalized utility of 0.52; 20 RFs, normalized utility ∼ 0, respectively) suggests that these cells might be a subject to additional unknown constraints or might be optimizing a different utility. We emphasize that we analyze the optimality of *individual* neurons, whereas the optimization of complete populations could yield a more diverse set of RFs that are individually suboptimally sparse *β*4, 38, 56), accounting for the deviations we observe. Our analysis is intended as a demonstration of the applicability of our framework, rather than a definitive optimality claim about V1 neurons. Population-level analysis of optimality is a subject of future work.

### Application 2: Receptive fields in the retina

Here we analyze temporal receptive fields of 117 retinal ganglion cells (RGCs) in the rat retina (57). Temporal RFs have a characteristic bimodal shape (Fig. 7A, left) which can be captured well by a simple filter model with three parameters (58). Two parameters (*c*_1_, *c*_2_) describe the amplitudes of both modes, while the third (*a*) determines the temporal scale of the filter (Fig. 7A, right panel). In what follows, we focus on the optimality of filter shapes in the space of these three parameters. RGC receptive fields long have been hypothesized to instantiate predictive coding (PC) – a canonical example of a normative theory in sensory neuroscience (59). Temporal PC postulates that, instead of tracking the exact stimulus value directly in their responses, neurons encode a difference between the stimulus and its linear prediction computed using past stimuli. Such a strategy has many potential benefits: it reduces the dynamic range of signals, minimizes use of metabolic resources, and can lead to efficient coding in the low noise limit, by performing stimulus decorrelation and response whitening (2, 55, 59–61).

**Fig. 7.**
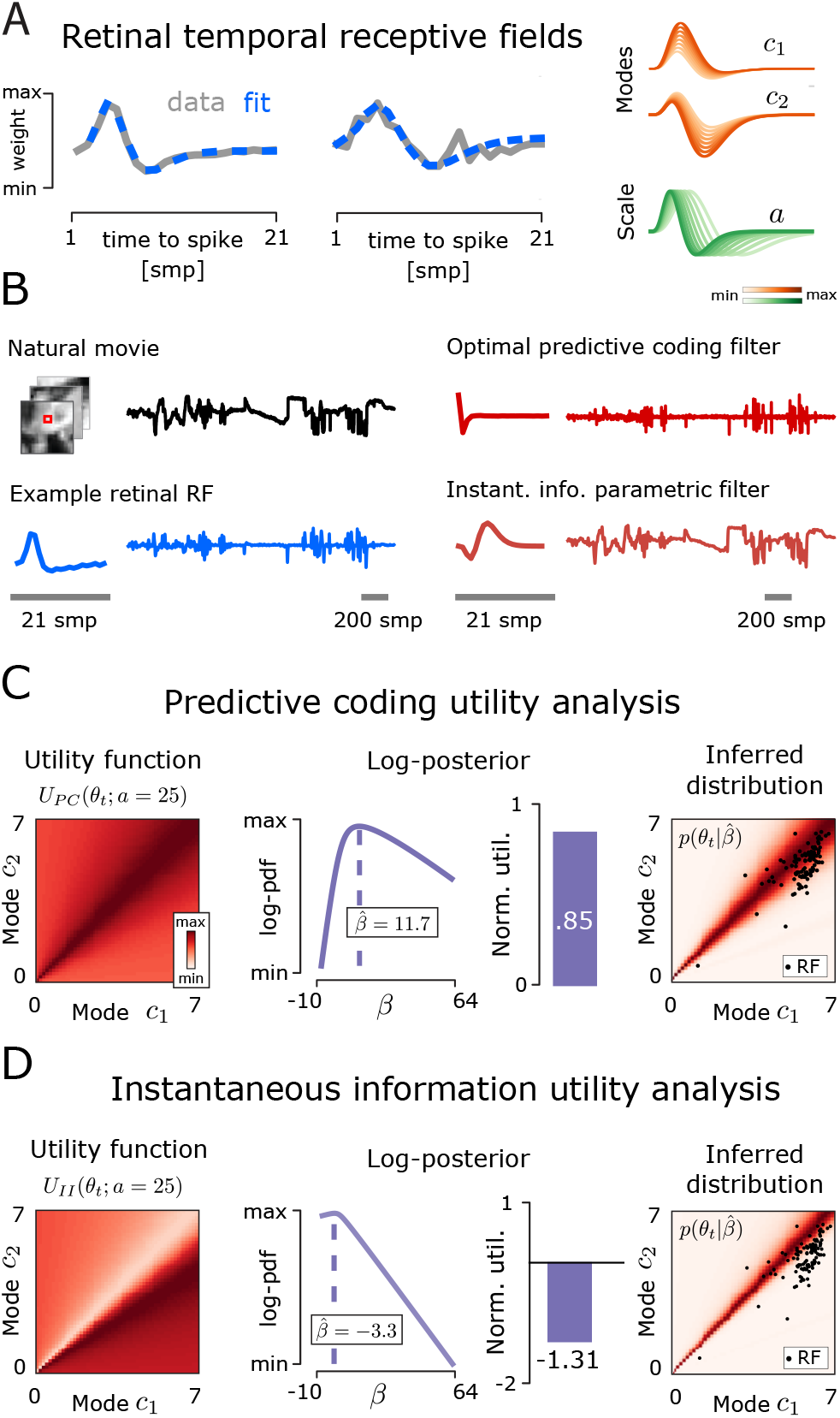
Optimality of retinal receptive fields. **(A)** Two example temporal receptive fields of rat retinal ganglion cells. Gray lines show RF estimates (courtesy of Olivier Marre (57)), dashed blue lines show parametric fits. Fit parameters correspond to amplitudes of filter modes (parameters *c*_1_, *c*_2_, orange) and scale (parameter *a*, green). **(B)** Example natural stimulus: light intensity of a single pixel of a natural movie (top-left, black). Representative retinal RF and its linear response to the natural stimulus (bottom-left, blue line). Optimal predictive coding filter and its response to the same stimulus (top-right, dark red line). Optimal instantaneous information transmission filter and its response (bottom-right, pink line). **(C)** Analysis of temporal RFs with the generalized predictive coding utility, *U*_PC_. First panel: Utility function of filter modes *c*_1_, *c*_2_ constrained by timescale *a* = 25. Second panel: Log-posterior (solid purple line) over population optimality parameter *β* (dashed vertical line – MAP estimate). Third panel: Normalized utility of the RF population. Fourth panel: Optimization prior distribution over (*c*_1_, *c*_2_) at the inferred 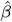, marginalized over all values of the timescale parameter *a* (black dots – data-derived RFs). **(D)** Analysis of temporal RFs with the instantaneous information utility, *U*_II_, analogous to C.

An optimal predictive coding filter must be adapted to the statistics of stimuli it encodes (59). We optimize PC filters using natural light intensity time-courses (see Methods). Optimal PC filter responses qualitatively resemble the responses of a representative retinal filter convolved with the same natural stimulus (Fig. 7C). Both filters generate strong, spike-like transients to sudden changes in the stimulus mean, while their output remains close to 0 when the stimulus is not changing. This pattern is different from the response of a parametric bimodal filter (with parameters *a, c*_1_, *c*_2_) optimized to track the stimulus, obtained by maximizing instantaneous information transmission in a low-noise regime (*U*_II_, see Methods). Importantly, predicted responses can be very distinct despite the qualitative similarity between retinal, PC, and instantaneous information filters.

To evaluate the optimality of retinal RFs, we propose a new utility, *U*_PC_, that mathematically generalizes the canonical formulation of predictive coding (59). This utility prioritizes filters which minimize power in their output, given a fixed filter norm, while allowing the filters to operate on timescales distinct from the stimulus frame rate (see Methods for details). We evaluate *U*_PC_(*θ*; *a*) as a function of the two filter mode parameters, *θ* = (*c*_1_, *c*_2_), but consider the timescale *a* to be an external constraint to be inferred from data for each neuron separately, as in Question 3. Parameter *a* is a constraint because, much like *k* in the toy neuron example of Fig. 2, its value is not set by optimality (which prefers *a* → 0) but by biophysical constraints or by the temporal horizon at which prediction is of highest use to the organism. For a broad range of *a* values, *U*_PC_ is highest close to the diagonal of the (*c*_1_, *c*_2_) plane, representing nearly balanced filters, as shown in Fig. 7C (left).

We use all retinal RFs jointly to compute the posterior over the optimality parameter *β* (Fig. 7C, second panel). The inferred 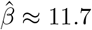 yields a normalized utility of 0.85, implying strong optimization for PC (Fig. 7C, third panel from the left); even relative to the non-parametric optimal PC filter with no timescale constraint, the utility of retinal filters remains as high as 0.74. The high degree of optimization is visually evident in the (*c*_1_, *c*_2_) plane, where individual neurons fall onto high utility regions of the maximum entropy distribution given inferred 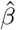 and marginalized over timescale *a* (Fig. 7C, right). An analogous analysis performed using maximization of instantaneous information *U*_II_ (see Methods) reveals a negative *β* estimate and thus anti-optimization for this alternative utility, with real neurons avoiding high-utility regions of the maximum entropy distribution.

### Application 3: Neural wiring in *C. elegans*

Here we analyze neural wiring in *C. elegans*, which has been the subject of several normative studies (20, 62–64). Relative positions of neurons could be partially predicted by minimizing the total wiring cost under the constraint that muscles and sensors need to be properly connected (62, 64). Instead of trying to predict individual neuron positions, we ask a different question: Are the measured neuron positions optimized to minimize the wiring cost to muscles and sensors?

For each neuron *i*, the wiring cost is determined by the number of muscles it connects to, the distance between the neuron’s position, *x*_*i*_, and positions of muscles, *m*_*i,j*_, and the number of synapses formed by each connection, *n*_*i,j*_ (Fig. 8A). The resulting utility function for each neuron can be written as 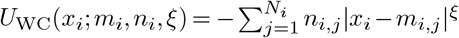, where *N*_*i*_ is the number of muscles the neuron *i* connects with, and *ξ* is an exponent determining the form of the utility as a function of distance (62) (Fig. 8B). The precise value of *ξ* is not specified by the theory and thus needs to be disambiguated from data, as described in Question 3.

**Fig. 8.**
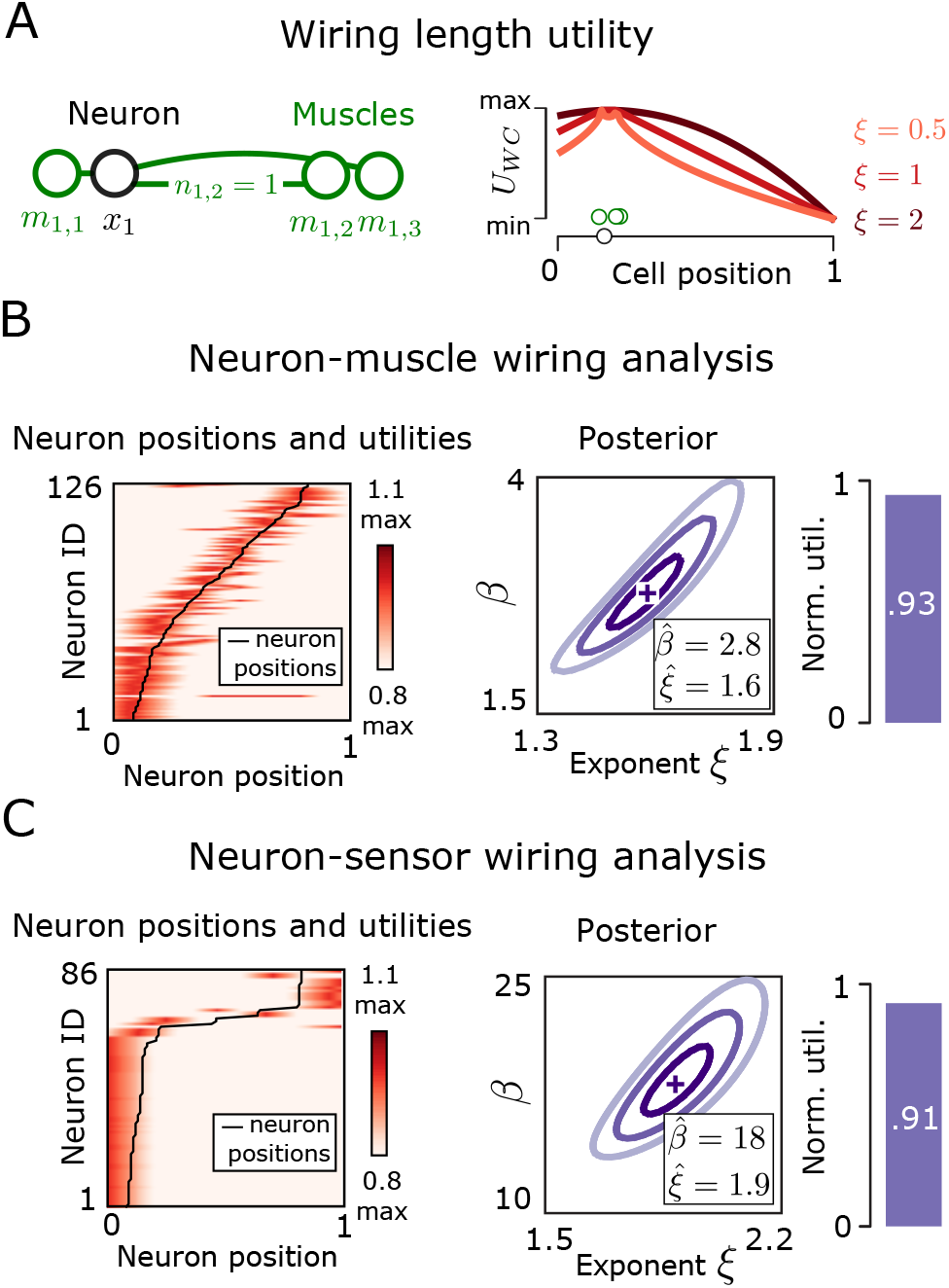
Optimality of neural wiring in *C. elegans*. **(A)** Left panel: Connection schematic between example neuron at position *x*_1_ (black circle) and three muscles at positions *m*_1,1_, *m*_1,2_, *m*_1,3_ (green circles). Number of synapses between neuron *x*_1_ and muscle *m*_1,*j*_ is denoted *n*_1,*j*_. The example neuron forms monosynaptic connections (green lines) only with the three muscles. Right panel: wiring cost utility, *U*_WC_(*x*_1_; *ξ*), as a function of position *x*_1_, corresponding to the scenario depicted at left. Position axis spans the entire *C. elegans* body length. Utility functions are shown for three exponent values *ξ*. **(B)** Neuron-muscle connection analysis. Left panel: Utility *U*_WC_(*x*; *ξ* = 2) (red, scaled to [0, 1] for each neuron) for all 126 neurons (rows), as a function of neuron positions *x* ∈ [0, 1]. Black line denotes positions of real neurons. Middle panel: joint posterior over optimality parameter *β* and the exponent *ξ* (cross denotes MAP estimates reported in the legend). Right panel: normalized utility of neuron-muscle connectivity. **(C)** Neuron-sensor connection analysis, analogous to B.

Our analysis shows that a large proportion of 126 neurons that form connections with muscles align closely with the maxima of the utility function (Fig. 8B, left panel). We estimate the joint posterior distribution over the optimality parameter *β* and the connection exponent *ξ*, for neuron-muscle and neuron-sensor connections separately (Figs. 8B,C, middle panels). In both cases, the normalized utility exceeds 90 %, implying strong optimization. Interestingly, the estimates for the exponent *ξ* are relatively high: 1.6 for neuron-muscle connections and 1.9 for neuron-sensor connections, suggesting that neurons are only weakly penalized for small deviations from optimal positions. This is in contrast to previously published analysis that focused instead on neuron-neuron connections (20), where the authors find (and we confirm) *ξ* ≈ 0.5. We interpret this discrepancy to imply that neuron-muscle and neuron-sensor connection costs are less important relative to neuron-neuron connections so far as the overall *C. elegans* body plan is concerned. One possible reason is that neuron-neuron wiring cost scales quadratically with the number of neurons, implying higher penalty (and thus lower *ξ*) for deviations from optimality.

## Discussion

In this paper we present a statistical framework that unifies normative, top-down models of complex systems which derive system parameters *ab initio* from an optimization principle, with bottom-up probabilistic models which fit system parameters to data. The union of these two approaches, often applied separately, becomes straightforward in the Bayesian framework, where the normative theory enters as the prior and data enters as the likelihood. The two traditional approaches are recovered as limiting cases; more importantly, interpolation between these two limits spans a mixed regime of optimization and inference that is highly relevant for understanding complex biological systems. We illustrated the relevance of our framework by describing how (i) measurements can be used to test a given system for consistency with a normative theory; (ii) “closeness to optimality” can be defined and inferred; (iii) ambiguities and degeneracies of theoretical predictions can be resolved, often by a small amount of data; (iv) normative theories can provide powerful priors to aid inference in high-dimensional problems.

Despite their theoretical appeal, the application of optimization principles to biological systems has been hindered by statistical issues that grow more pressing as the complexity and dimensionality of the models increases. These issues are not new. Instead of developing an *ad hoc* solution whenever called for by a particular application, we decided to tackle these issues head on and flesh them out with simple examples. For instance, the issue of an unconstrained optimization parameter or a trade-off with unknown strength is well-known to the practitioners, but is often solved “by hand”: one manually adjusts the constraint until the optimality predictions are (visually) consistent with data. Such manual “fine-tuning” of constraints is clearly problematic from the statistical viewpoint, as it could easily amount to (over-)fitting that is not controlled for. In contrast, our framework performs inference and optimization jointly and provides a full posterior over constrained and unconstrained parameters alike. Another problematic issue arises from degenerate maxima of the utility functions. A frequent solution has been to postulate further constraints within the theory itself, which disambiguate the predictions (15). Our framework proposes a complementary mechanism: using a small amount of data to localize the theoretical predictions to the relevant optimum, against which further statistical tests can be carried out. As a last example, when fitting complex (e.g., nonlinear dynamical systems) models one typically restricts parameters by hand to a domain that is thought to be “biologically relevant.” In contrast, optimization priors automatically suppress vast swaths of parameter space that lead to non-functioning systems, even if these systems are not fully optimized for the postulated utility. In this way, the statistical power of the data can be used with maximum effect in the parameter regime that is of actual biological relevance, without sacrificing statistical rigor.

While our framework provides a principled way to navigate these and similar statistical issues, important questions remain. A key challenge is to identify the relevant optimization criterion for a biological system, and to express it in terms of experimentally measurable quantities. A candidate utility function which embodies an optimality criterion of interest could be selected from a possible discrete set of such functions (17, 61, 65), or by inferring utility function parameters. Because we leverage the well-understood machinery of Bayesian inference, one could perform model selection for the utility function that best explains the data. Such an approach could be used, for example, to rigorously verify whether entire neural populations in the visual cortex are jointly optimized for sparsity or a different utility, such as slowness (54). An important caveat is that the more flexible our choice of the utility function becomes, the easier it is to claim an optimality for a system of interest. In principle, one could postulate a utility function with a fully unconstrained shape: in this limit, our framework would automatically recover the utility function shape from data (if these were sufficient) assuming the observed system is optimal, in a way reminiscent of inverse reinforcement learning (61). This connection is an interesting topic for further research. In this paper, however, we focused on optimization theories where the number of adjustable utility parameters is smaller than the number of system parameters being predicted.

Our framework dovetails with other approaches which address the issues of ambiguity of theoretical predictions and model identifiability given limited data in biology. “Sloppy-modelling” (66, 67), grounded in dynamical systems theory, characterizes the dimensions of the parameter space which yield qualitatively similar behavior of the system. In our framework, these dimensions correspond to regions of the parameter space of equal or similar utility. Another important conceptual advance grounded in statistical inference has been the usage of limited data to coarse-grain probabilistic models (26, 68, 69). In our framework, a related coarse-graining occurs when, instead of inferring all system parameters from data directly, optimization sets the values of most of these parameters, leaving only the unconstrained subset to be fitted. The resulting dimensionality reduction could be sizable (e.g., with optimization predicting high-dimensional RF shapes given inferred firing rate, locality, or neural noise constraints), and could efficiently parametrize neuronal heterogeneity in terms of a small number of constraints that vary from neuron to neuron or between neural populations. Another point of connection with recent work concerns the ability to instantiate high-dimensional maximum entropy distributions over parameters with complicated dependency structures (16, 21, 70). Such computational innovations will be essential for statistical analyses of optimality that require sampling from maximum-entropy optimization priors.

Theories of biological function are currently less structured than physical theories of nonliving matter. This is partially due to the inherent properties of biological systems such as intrinsic complexity and lack of clear symmetries. It is also partially due to the lack of theoretical approaches to systematically coarse-grain across scales and identify relevant parameters. We hope that our approach which synthesizes statistical physics, inference, and optimality theories, can provide novel ways in tackling these fundamental issues.

## Acknowledgements

The authors thank Dario Ringach for providing V1 receptive fields and Olivier Marre for providing retinal receptive fields.

## Methods

### Key resources table

#### Resource availability

Further information and requests for code should be directed to Wiktor Młynarski. This study did not generate any new datasets. The code supporting the current study is freely available from the corresponding author upon request.

### Method details

#### Model neuron and mutual information utility function

A model neuron elicits a spike at time *t* (*r*_*t*_ = 1) with a probability:

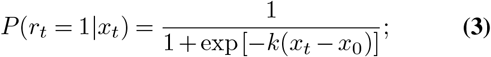

the stimuli *x*_*t*_ were distributed according to a Gaussian Mixture Model, 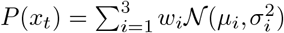, where *w*_*i*_ = 1*β* are weights of the mixture components, *µ*_1,2,3_ = 2, 0, 2 are the means, and *‡*_*i*_ = 0.2 are standard deviations.

To estimate mutual information between class labels and neural responses, we generated 5·10^4^ stimulus samples *x*_*t*_ from the stimulus distribution. Each sample was associated with a class label *c*_*t*_ ∈ {1, 2, 3 }, corresponding to a mixture component. We created a discrete grid of logistic-nonlinearity parameters by uniformly discretizing ranges of slope *k* ∈ [−10, 10] and position *x*_0_ ∈ [− 3, 3] into 128 values each. For each pair of parameters on the grid, we simulated responses of the model neuron to the stimulus dataset and estimated the mutual information directly from a joint histogram of responses *r*_*t*_ and class labels *c*_*t*_.

### Likelihood ratio test of optimality

The proposed test uses the likelihood ratio statistic,

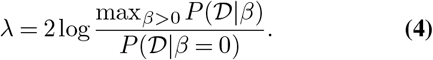

The null hypothesis is rejected for high values of *λ*. The marginal likelihood of *β*, 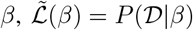, depends on the overlap of parameter likelihood and the optimization prior, 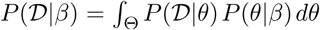, where ⊝ is the region of biophysically feasible parameter combinations.

The null distribution of *λ* is obtained by sampling in three steps: (i) sample a parameter combination *θ* from a uniform distribution on *θ*, i.e. *P* (*θ*|*β* = 0); (ii) sample a data set 𝒟 according to the likelihood *P* (𝒟|*θ*); (iii) compute the test statistic *λ* according to Eq. (4). This computationally expensive process simplifies in two situations described below.

### Data-rich-regime simplification

In the data-rich regime, when the parameter likelihood *P* (𝒟|*θ*) is concentrated at a sharp peak positioned at 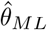, likelihood ratio depends only on the value of utility at 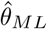:

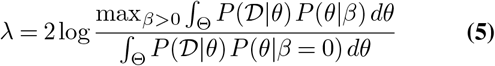

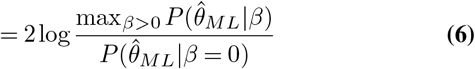

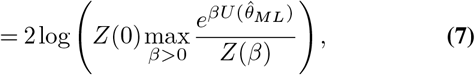

which is a non-decreasing function of the utility 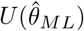. Thus, this test is equivalent to a test that uses the utility estimate itself, 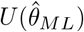, as the test statistic, making it possible to avoid the costly integration over ⊝. The null distribution can then be obtained by computing *U* (*θ*) at uniformly sampled *θ*.

### Multiple system instances simplification

If multiple instances of the system are available and we can assume that their parameters *θ*_1_, *θ*_2_,…, *θ*_*N*_ are i.i.d. samples from the same distribution *P* (*θ*|*β*), then the datasets 𝒟_1_, 𝒟_2_,…, 𝒟_*N*_ are also i.i.d., 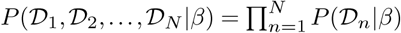. We test the hypotheses *β* =0 vs. *β >* 0 with the likelihood ratio statistic

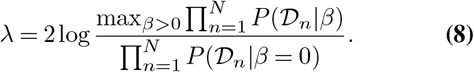

By Wilks’ theorem, for large *N* the null distribution of *λ* approaches the 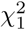 distribution (with a point mass of weight 1*/*2 at *λ* = 0, because we only consider *β* ≥0). This removes the need for sampling in order to obtain the null distribution.

#### Hierarchical inference of population optimality

Assuming that experimental datasets 𝒟_1_, 𝒟_2_,…, 𝒟_*N*_ are i.i.d., the posterior over population optimality parameter *β* takes the form:

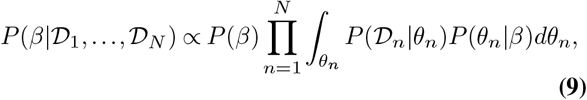

where *θ* = (*k*_*n*_, *x*_0,*n*_) is a vector of neural parameters (slope and position), and *P* (*β*) is a prior over *β*. We approximated integrals numerically via the method of squares. Neural parameter values were sampled from ground-truth distributions via rejection sampling.

#### Inference of receptive fields with optimality priors

We randomly sampled 16 × 16 pixel image patches from the van Hateren natural image database (55) and standardized them to zero mean and unit standard deviation. Neural responses were simulated using a Linear-Nonlinear Poisson (LNP) model:

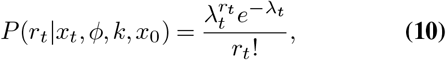

where *λ*_*t*_ is the rate parameter equal to:

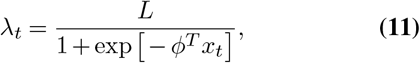

where *L* = 20 was the maximal firing rate.

Given a linear filter *ϕ*, we quantified sparsity of its responses to natural images using the following function:

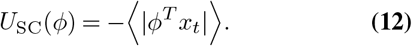

Filter sparsity was averaged across the natural image dataset consisting of 5*·*10^4^ standardized image patches randomly drawn from the van Hateren image database. The mean and standard deviation of filters *ϕ* was set to be 0 and 1 respectively. We optimized filters which either maximize or minimize the sparse utility measure via gradient descent. Different random initializations led to different filter shapes.

The locality utility of neural filters was defined as follows:

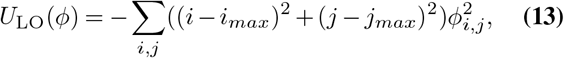

where *i*_*max*_, *j*_*max*_ are positions of the RF pixel with the largest absolute value. This definition of locality was introduced in (15).

Sparsity and locality utilities were combined into a single utility:

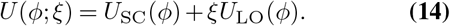

To estimate receptive fields (neural filters), we first simulated the responses of the model population to 2000 natural image patches. We estimated linear receptive fields from simulated data by computing the spike-triggered average (STA), a widely applied estimator of neural receptive fields (51). In the STA model, response of neuron *n* at time *t* is assumed to follow the normal distribution (18):

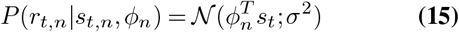

where *ϕ*_*n*_ is the linear receptive field of the n-th neuron, and *‡*^2^ is the noise variance.

To infer the receptive fields from simulated neural responses using our framework, we assumed the following optimization prior over receptive fields derived from the sparsity utility in Eq (12):

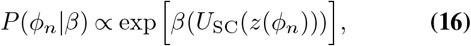

where *z*(*ϕ*_*n*_) denotes normalization of the receptive field to zero mean and unit variance. The sparse utility was evaluated over 10^4^ randomly sampled image patches. The resulting log-posterior took the following form:

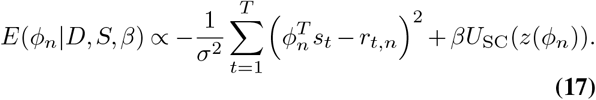

MAP inference was performed via gradient ascent on the log-posterior. Receptive fields were inferred with different priors corresponding to following values of the *β* parameter: 0, 1, 10, 20, 100. Receptive fields were estimated after reducing the dimensionality of stimuli with Principal Component Analysis to 64 dimensions. Estimation via gradient ascent on the log-posterior was performed in the PCA domain. PCA preprocessing is equivalent to low-pass filtering the stimuli. To estimate value of the locality constraint *ξ* as well as the prior strength *β* via cross-validation, we split the data into the training and testing datasets comprising of 80% and 20% of data respectively. We estimated receptive fields for a range of *β* and *ξ* values ([0, 0.01, 0.1, 1, 10] and [0, 0.05, 0.2, 1] respectively). For each MAP RF estimate, we predicted neural responses 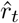 using stimuli from the test dataset. We then computed the average error 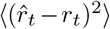 using neural responses in the test dataset. Combination of hyperparameters *ξ,β* which resulted in the smallest error value was taken to be the estimate of the correct one.

#### Analysis of V1 receptive fields

Receptive fields of 250 neurons in the Macaque V1 were published and analyzed in (49). All receptive fields were downsampled to 32 × 32 pixels size and normalized to have zero mean and unit variance.

To evaluate sparseness of V1 receptive fields, we relied on the following sparse utility:

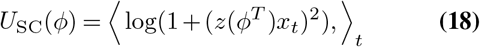

where *x*_*t*_ are individual image patches and *z*(*ϕ*_*n*_) denotes normalization of the receptive field to zero mean and unit variance. The sparse utility was evaluated over 5 × 10^4^ randomly sampled image patches. This form of the sparse utility was proposed in (71), and together with the measure specified in Eq (12) it belongs to a broad class of equivalent sparsity measures defined by convex functions (38).

To test individual RFs for optimality, we generated the null distribution of utility values by bootstrapping 10^6^ random filters as follows: (i) draw a random integer *K* between 1 and 128; (ii) superimpose *K* randomly selected principal components of natural image patches; each component is multiplied by a random coefficient *v* ϵ 𝒩; (0, 1); (iii) generate a 2D Gaussian spatial mask centered at a random position on the image patch; lengths of horizontal and vertical axes of the Gaussian ellipse were drawn independently; (iv) multiply the random filter and the Gaussian mask. This procedure ensures that a range of filters of different sparsity and slowness will be randomly generated. Filters were standardized to zero mean and unit standard deviation.

To establish a measure of optimality at a population level, we needed to simplify the integration over all receptive field parameters, which was intractable due to their high-dimensionality. Computation of posteriors over *β* in Eq (9) was therefore approximated as follows:

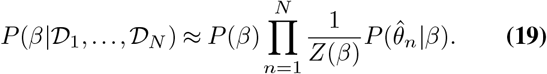

where 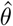 are receptive fields estimates computed in (49).

We approximated 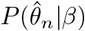 via rejection sampling, noting that 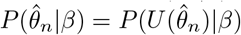, i.e., the probability of a high dimensional receptive field is determined solely by a one-dimensional utility function.

For each *β* we randomly sampled 10^6^ filters from the proposal distribution, as described above, and retained only those consistent with *P* (*U*_SC_(*θ*)|*β*) via rejection sampling. Obtained utility values were fitted with a Gaussian distribution, used to evaluate posteriors over *β*, with point estimates being posterior maxima; the prior over *β* was uniform over the range displayed in the figures. For sparse utility, we discretized *β* values into 20 values equally spaced on the [-5, 5] interval. For slow utility we used 64 *β* values equally spaced on the [32, 32] interval.

Filters accepted for each *β* value were used to compute the average spatial autocorrelation.

To cluster receptive fields according to optimality, we defined a mixture model:

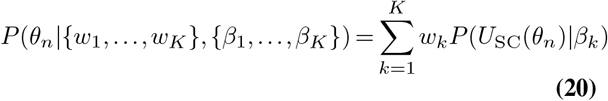

where *w*_*k*_ is the weight of the *k*th mixture component and *β*_*k*_ is the optimality of that component. To approximate utility-defined distributions, we used the Gaussian approximation described above i.e.: 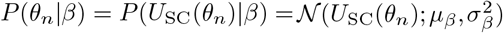

Parameters of the model were learned via the standard expectation-maximization algorithm (EM).

#### Analysis of retinal receptive fields

Temporal receptive fields of retinal ganglion cells were published and analyzed in (57). We analyzed RFs of 117 neurons selected by temporal smoothness. Each RF was normalized to unit norm and fitted with a parameteric biphasic filter model described in (58).

We considered two different utility functions. First one was a generalization of the predictive coding objective introduced in (59). The predictive coding objective minimizes the squared difference between the stimulus value *s*_*t*_ at time *t* and the linear prediction of that stimulus value computed from *N* past values: 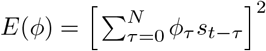, where *ϕ* are the weights of the linear filter. In the classical approach it has been assumed that the linear weight of the current stimulus *s*_*t*_ is equal to 1 i.e. *ϕ*_0_ = 1. We note that such form makes it difficult to evaluate predictive coding filters adapted to stimuli of unknown temporal scale. In particular, we optimize and evaluate our filters on natural movies whose frame rate might be mismatched with the timescale of the retina. We therefore relax the assumption that the predictive coding filter reduces the dynamic range by subtracting only the current stimulus from its prediction, and assume that what is being predicted is itself a linear combination of stimulus values (e.g., integrating stimulus value over some recent period of time). In practice this means that we allow all values of the filter including *ϕ*_0_ to vary freely. To avoid trivial solutions, where the residue *E*(*ϕ*) is minimized by setting all weights to 0, we impose a unit norm constraint on the filter *ϕ*. The utility function of a filter *ϕ* is then equal to:

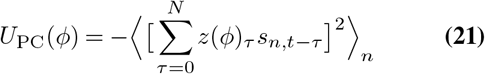

where *z* denotes the unit norm operator, and *n* indexes stimulus epochs *s*_*n*_.

We evaluated the utility *U*_PC_ using 50000, 21-sample long excerpts of single-pixel luminance extracted from natural movies of scenes in the African savanna (72).

We used these natural stimulus data to learn the optimal predictive-coding filter, as described in (59) via gradient descent.

The second considered utility was measuring the amount of information between the stimulus and the instantaneous filter output in a low-noise regime. Under the Gaussian approximation of stimulus and output distribution this utility takes the form:

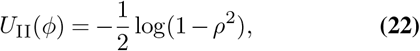

where *ρ* is the Pearson correlation coefficient between the stimulus *s*_*t*_ and the filter output *r*_*t*_. This utility is high when the neural responses track the stimulus with high fidelity. Note that this is not the general solution to an efficient coding (infomax) problem, where the *full response trajectory*, not the instantaneous response, should encode high information about the stimulus, which leads to decorrelation / whitening in the low-noise regime. We evaluated *U*_II_ using a trajectory of 20000 samples of pixel intensity values extracted from the natural movie dataset.

To compute utility-defined distributions of the filter mode amplitude parameters *c*_1_, *c*_2_, we first discretized values of these parameters into 100 values uniformly spaced on the [0.01, 13] interval, where 13 was the maximum amplitude parameter value among fits to normalized retinal RFs. For each filter we evaluated utility for each pair of discretized amplitude parameter values and a fixed value of the scale parameter *a* fitted to that filter. We used such utility surfaces to estimate the normalization constant of the utility-defined distribution parametrized by *β* and the scale parameter *a*.

We discretized the parameter *β* into 100 values uniformly spaced on the [-10, 64] interval. We estimated the posterior over *β* by numerically integrating over filter parameters *c*_1_, *c*_2_, *a*. We assumed a uniform prior over *β*.

#### *Analysis of connectivity in* C. elegans

For our analysis we used the *C. elegans* neural wiring dataset available on Worm Atlas (www.wormatlas.org). This dataset has been published and analyzed before in (62) as well as (20, 64) – for details about the dataset please refer to this prior work.

For the analyses depicted in Fig. 8 we selected two sets of neurons. The first set consisted of 126 neurons connected to at least one muscle, and the second set consisted of 86 neurons connected to at least one sensor. “i-th” neuron was therefore characterized by its position, *x*_*i*_, number of land-mark cells (muscles or sensors) it was connected to, *N*_*i*_, vectors of positions of the landmark cells, *m*_*i*_ (muscles), and *s*_*i*_ (sensors), and vectors of the number of synapses in each neuron-to-landmark connection, *n*_*i*_. For each neuron the utility of its position was defined as:

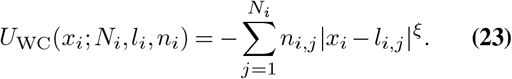

where *l œ* {*m*_*i*_, *s*_*i*_ }, denotes the vector of landmark cell positions. We evaluated the utility function on the [0, 1] interval representing the linear extent of the worm body axis, discretized into 100 linearly spaced values. To compute the posterior distribution over parameters *β* and *ξ* we discretized them into 64 linearly spaced values. For neuron-muscle connections, *β* was defined over a [1.5, 4] interval and *ξ* over a [1.3, 1.9] interval. For neuron-sensor connections, *β* was defined over a [10, 25] interval and *ξ* over a [1.5, 2.2] interval. We assumed a uniform prior over parameters *β, ξ*.

### Quantification and statistical analysis

Statistical test performed in Fig. 5D was a two-tailed t-test. Stars denote p-values lower than 0.001. Error bars in the figure denote standard errors of the mean.

## S1 Effects of parametrization

Biological systems can often be described using different sets of parameters. One set of parameters can be converted into another using a mathematical transformation, such as the log-transform (replacing the value of slope *k* by log *k*) or the reciprocal (replacing *k* by 1*/k*). The choice of parametrization is largely a matter of convenience or convention. However, when working with probability distribution functions such as the optimization priors (Eq. 2), special care needs to be taken, since the functional form of the priors is intertwined with the choice of parametrization. In this section we highlight the subtleties involved. In the next section S2 we introduce a generalized form of the optimization priors, that allows us to clearly separate the choice of the parametrization from the choice of the normative prior distribution.

### Maximum entropy priors depend on parametrization

Suppose that a researcher, Alice, defines the family of optimization priors as introduced in Eq. 2,

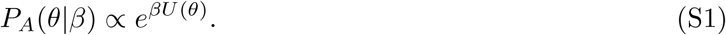

Suppose that Alice then decides to use a different set of parameters, *ϕ* = *f* (*θ*), to perform some computational task. The prior (S1), as function of *ϕ*, can be obtained by the standard method of changing variables,

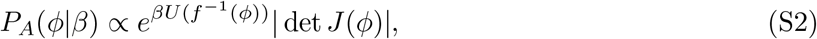

where 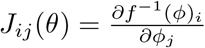 is the Jacobian. Some parameter regions may appear expanded or shrunk in the new parametrization; the Jacobian corrects for this and makes sure that the underlying probability distribution does not change. A change of variables like this can be done whenever it is convenient.

Notice, however, that the distribution *P*_*A*_(*ϕ*|*β*) in (S2) is different from what another researcher, Bob, who has been using the parametrization *ϕ* from the beginning, has obtained from Eq. 2 directly,

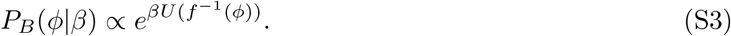

Namely, Bob’s distribution *P*_*B*_(*ϕ*| *β*) does not include the Jacobian that is present in *P*_*A*_(*ϕ*|*β*), since Bob did not perform a change of variables from *θ*. This means that Alice and Bob are using different optimization priors and will get different results downstream in the analysis.

This is particularly clear for *β* = 0, i.e. with zero optimization. Bob’s prior *P*_*B*_(*ϕ*|*β*) is then uniform in *ϕ*; but Alice’s prior *P*_*A*_(*ϕ*|*β*) ∝ | det *J*(*ϕ*)| is uniform in *θ*, but non-uniform in *ϕ* (unless *θ* and *ϕ* are related by a linear transformation, in which case the Jacobian | det *J*(*ϕ*)| is constant).

### Illustration with the toy model

Fig. S1 shows three different parametrizations of our toy model,

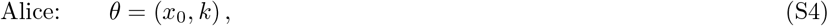

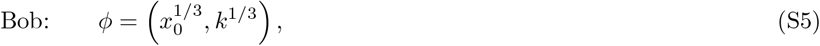

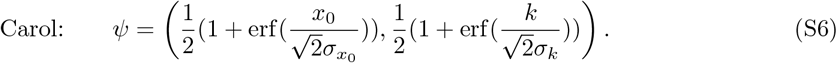

The ψ parametrization corresponds to the Gaussian distribution that we used as a non-optimised example in the paper, Fig. 4C (III).

In Fig. S1, column A shows the utility surface in the three different parametrizations. Compared to Alice’s parametrization *θ* (top row; this is the parametrization also used throughout the paper), the parameter regions around (*x*_0_, *k*) = (0, 0) are inflated in Bob’s and Carol’s parametrizations *ϕ* and (middle and bottom rows). Columns B-D show the likelihood surfaces, which show the same distortion between the parametrizations. Column E in Fig. S1 shows the optimization priors of Alice, Bob and Carol under no optimization, *β* = 0, transformed into Alice’s parameters *θ*. Alice has defined her prior family according to Eq. 2 using *θ* and it is therefore uniform for *β* = 0. Bob and Carol have defined their priors to be uniform *ϕ* and, and they are therefore not uniform in *θ*. Specifically, the region around (*x*_0_, *k*) = (0, 0) which was inflated in *ϕ* and has higher probability density when “shrunk” back to *θ*. We later refer to the *β* = 0 optimization prior, *P* (*θ*|*β* = 0), as the *null model*.

Since Alice, Bob and Carol are effectively using different priors, they obtain different results in downstream analyses. To demonstrate this, Fig. S2 shows the likelihood of *β* for the three example systems (subplots) and three parametrizations (blue, orange, green). The differences between parametrizations are mostly qualitative for examples 1 and 3, which are on the two extremes – non-optimised and highly optimised (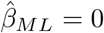and, ∞ regardless of parametrization).

In the intermediate example 2, the ML estimates vary based on parametrization. For example,squeezes the utility peaks into the corners (Fig. S1A, bottom). This leads to 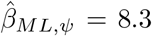, higher than 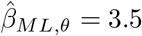 for the original parametrization, where the peaks are more spread out.

These differences raise the question of which parametrization leads to correct results. First, we stress that the answer is problem-specific and cannot be addressed in general. One should consider whether a prior uniform in the chosen parameters under zero optimization (*β* = 0) is appropriate.

Second, whenever this is desirable, the choice of parameters can be decoupled from the prior distribution. A change in variables can be performed after the optimization priors are defined. Alternatively, a null distribution under zero optimization can be specified explicitly – this is discussed in the following section.

## S2 Optimization priors with general null models

The maximum entropy optimization priors from Eq. 2 are uniform for *β* = 0 (we refer to the *P* (*θ*|*β* = 0) as the *null model*). However, finding a parametrization where this is appropriate (see previous section S1) can be difficult. For example, how to parametrize receptive fields such that a reasonable null model is uniform in some domain? The choice of parameters can also be dictated by convenience or convention. For such cases and in general, we can decouple the choice of parametrization from the choice of the null model.

**Figure S1:**
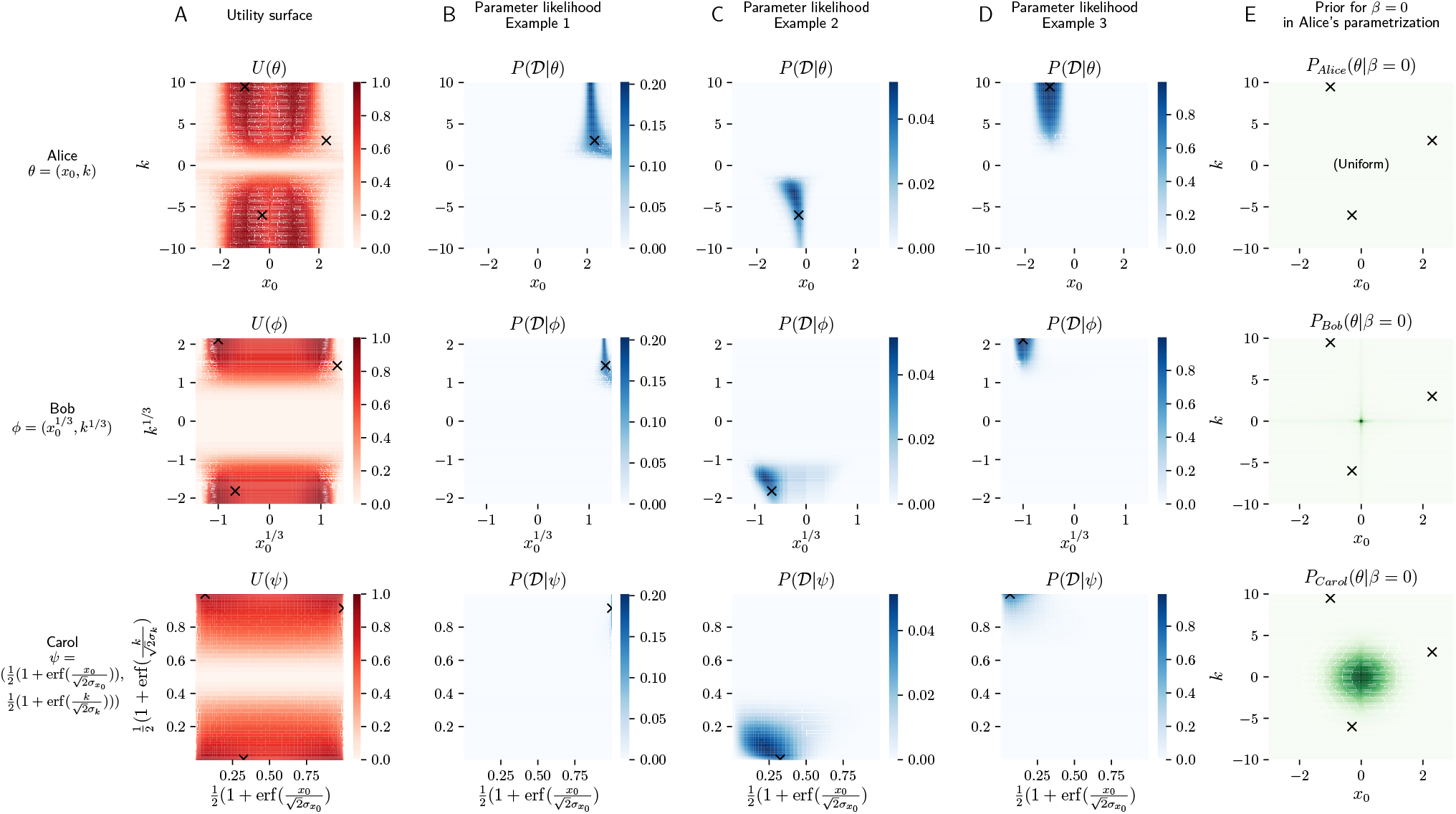
An illustration three different parametrizations of the toy model. Alice, Bob and Carol use *θ, cf* and – shown in top, middle and bottom row respectively. Column **A**: Utility plotted in the different parametrizations. Columns **B-D**: Likelihood from the three examples used in the main text Fig. 3B, here plotted in 3 different parametrizations. Column **E**: Optimization priors under *β* = 0 that Alice, Bob and Carol are effectively using when they applied Eq. 2 using their choice of parametrization. For comparison, all are plotted in Alice’s parametrization *θ*.

**Figure S2:**
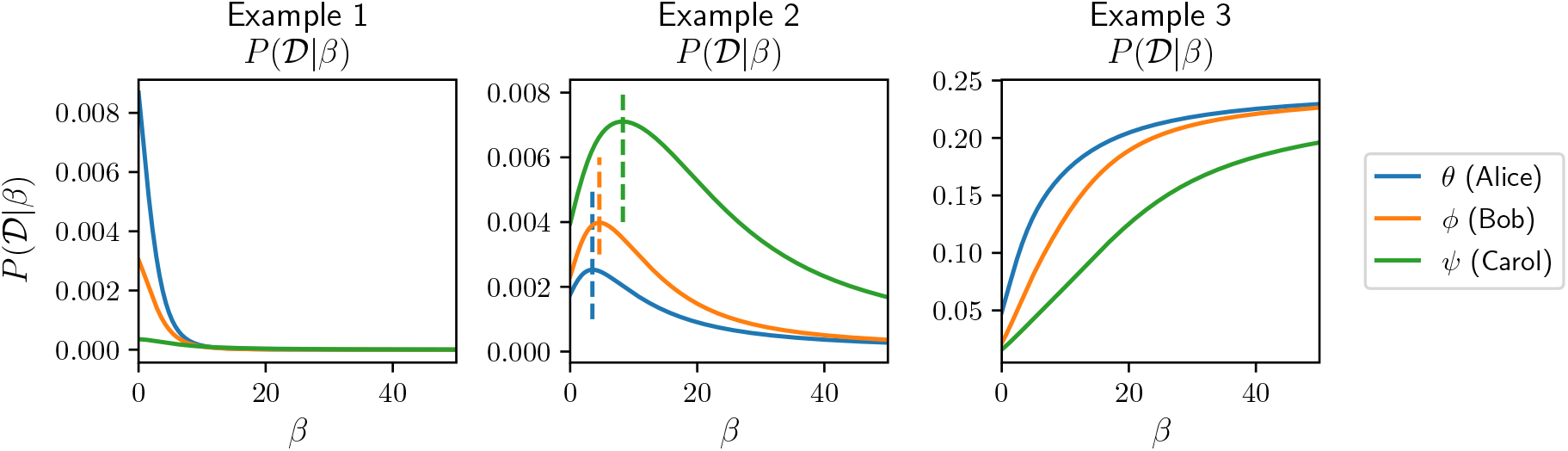
Likelihood of *β* for three example systems and three parametrizations *θ, cf* and. **Example 1** (see likelihood in Fig. S1B) is not optimised and ML *β* is always 0, but the curve differs between parametrizations. Similarly, **Example 3** (likelihood in Fig. S1D) is strongly optimised and ML *β* is always *1*. The intermediate **Example 2** (likelihood in Fig. S1C) has finite ML estimate of *β*, which depends on parametrization (dashed vertical lines). The values are 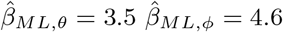 and 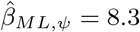.

Let the null model – the parameter distribution without optimization – be *q*(*θ*). Then the normative prior family

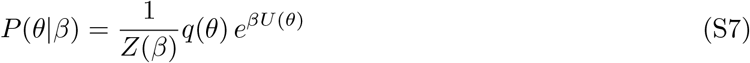

is maximum entropy in the sense that is solves the optimization problem

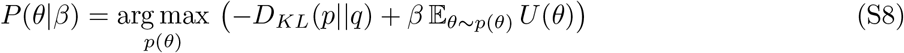

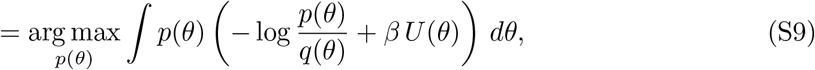

where *D*_*KL*_ is the Kullback-Leibler divergence or relative entropy. Intuitively, *P* (*θ*|*β*) is as similar to *q*(*θ*) as possible, while constraining average utility. If *q*(*θ*) is uniform in some domain, we recover the maximum entropy solution from Eq. 2.

When changing parametrization, *q* will transform accordingly. E.g. Alice and Bob might choose two null models *q*_*A*_(*θ*) and *q*_*B*_(*ϕ*), each using their preferred parametrization. They are equivalent (Alice and Bob will get the same results) if

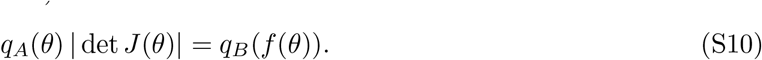

### How to choose the right parametrization/null model?

In general, the choice is problem-specific and beyond the scope of our paper. Note that even when *q*(*θ*) is uniform on some domain, the domain needs to be chosen and affects the results.

In some biological systems, an established null model may be available. Distributions of the form (S7) have a history in population genetics [1], [2] of describing the equilibrium distributions of allele frequencies; the functional form emerges from a stochastic model of evolution. The null model *q*(*θ*) is then the equilibrium distribution under mutation and random drift; natural selection enters through the factor *e*^*βU*(*θ*)^. The parameter *β* quantifies the strength of natural selection in favour of *U* relative to random drift.

In other cases, the null model may simply reflect our limited knowledge about the system. As a half-joke example, a simple null model for positions of neurons along the AP axis in *C. elegans* might be uniform, ignoring the intricacies of its body shape. The situation is different for an animal like the brontosaurus, since “All brontosauruses are thin at one end, much, much thicker in the middle, and then thin again at the far end” [3]. More elaborate null models might take into account cell lineages, or assume that neurons are allowed to permute their positions, but not occupy new places (making *θ* discrete).

In the example of visual receptive fields, it may be worth considering if properties such as smoothness should be included in the null model *q* or in the utility part of the prior *e*^*βU*(*θ*)^. If smoothness is present without optimization, then for the purposes of testing and quantifying optimality, it should be included in *q*. For Bayesian inference this might not matter, since performance is more important than the justification of the prior/null model.

## S3 Interpreting the magnitude of optimization parameter *β*

The optimization parameter *β* enters the optimization priors in a product with utility, *e*^*βU*(*θ*)^. This means that the magnitude of *β* cannot be directly compared between different biological systems, or different utility functions considered in the same system. However, there are several ways to interpret the size of *β*.

### Probability of observing a parameter combination

As follows from the functional form of the optimization priors, each additional unit of utility *U* makes a parameter combination *θ* more probable by a factor *e*^*β*^. One can ask how much of a probability increase is then conferred by a biologically significant increase in *U*.

### Normalized utility

Each value of *β* has an associated average utility 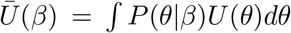 achieved by the optimization prior. Normalized utility,

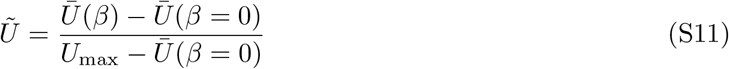

is plotted throughout the paper. This converts *β* to the expected, biologically meaningful increase in utility, and where it falls on the scale from not optimized at all (at *β* = 0) and maximum possible utility (*U*_max_). Negative values for normalized utility are possible for cases of “anti-optimization”, when the inferred *β <* 0.

Note that normalized utility is a distributional property, since it is defined as an average value of utility over an optimization prior at an inferred value of *β*. It is also possible to evaluate and normalize the point estimate of utility at a particular value of (e.g. inferred) parameters *θ*, 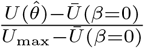. This may be more appropriate if we only deal with data for a single instance of a system, which may not permit a reliable estimate of *β*.

The mapping between *β* and *Ū* (*β*) is not trivial and depends on the distribution of *U* under no optimization. A direct calculation shows that the average utility grows with its variance,

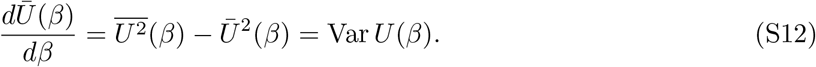

In particular, close to *β* = 0, *Ū* (*β*) grows linearly with slope Var *U* (*β* = 0). This can be expected intuitively – if the available parameter combinations provide a large variety of utility values, even small *β* (weak selection, see below) can induce a large change in *Ū*.

If the distribution of utility under *β* = 0 is Gaussian, this linear growth continues indefinitely, otherwise the growth is nonlinear and depends on higher moments of that distribution.

### Alternative normalization of utility functions

Based on the above arguments, it would be also possible to “standardize” the raw utility values by subtracting the average utility at *β* = 0 and dividing by Std *U* (*β* = 0). Such standardized utility would then enter the optimization prior and inference. In this representation, inferred *β* values would be directly interpretable: an increase of 1 for the value of *β* would lead to an approximate increase of 1 in the standardized utility; while zero, as before, would correspond to the expected utility with no optimization. The advantage of this method is that *U*_max_ need not be known in advance (which could be intractable to find exactly in high-dimensional spaces); the drawback is that one needs to estimate *Ū* (*β* = 0) and Std *U* (*β* = 0) by Monte Carlo sampling to standardize the utility *before* doing any inference. We did not use this alternative normalization in the paper.

### Strength of natural selection

A factor analogous to *e*^*βU*(*θ*)^ appears in the equilibrium distribution of allele frequencies [1], [2] in population genetics, if log fitness if proportional to *U*. The parameter *β* corresponds to the log fitness advantage per unit utility, multiplied by the effective population size. Fitness is the expected number of offspring in the next generation; population size enters because selection is more efficient in larger populations, where stochastic effects (random drift) are weaker.

## S4 Trade-offs between utility and likelihood

The normative priors defined in Eq. (2), and the optimization parameter *β*, are used in a Bayesian framework to interpolate between predictions from normative theories and inferences from data. Along this interpolation path, various trade-offs between theory and data are made. Here we focus on the maximum posterior parameter estimates, and show that they achieve the best possible utility-likelihood trade off. This provides an additional justification for the maximum entropy form of the optimization priors, Eq. (2).

Consider the the maximum posterior (MP) estimates of *θ*, as a function of the optimization parameter *β*,

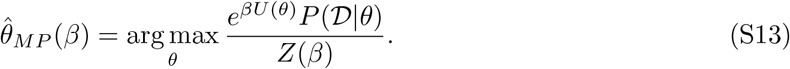

As *β* increases from 0 to infinity, 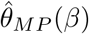 goes from the maximum likelihood estimate 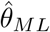 to the maximum utility prediction, *θ*^∗^.

The maximum posterior trajectory 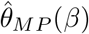, parametrized by *β*, achieves the best possible trade-off between the normative theory and data in the following sense: it finds the parameter combinations with the highest utility given each value of likelihood. Or conversely, it finds the most likely parameter combination, for a given each value of utility. This can be shown by rewriting the MP formula^1^ as

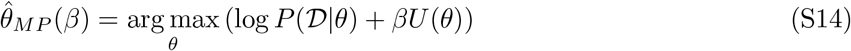

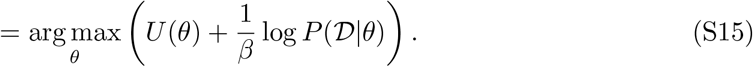

Here, *β* takes the role of a Lagrange multiplier constraining *U* (*θ*) when maximizing log *P* (𝒟|*θ*); or equivalently, 1*/β* constrains log *P* (𝒟|*θ*) when maximizing *U* (*θ*).

### Comparison with alternative interpolation methods

We compare the utility-likelihood trade-off achieved achieved by maximum posterior estimates based on the maximum entropy optimization priors with two alternative methods of interpolation between theory and data.

#### Linear interpolation between maximum likelihood and maximum utility parameters

In this approach we reduce both the normative theory and the data to points in the parameter space, maximum utility *θ*^∗^ and maximum likelihood 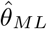, and interpolate between them, using

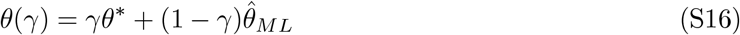

with higher values of parameter *γ* ∈ (0, 1) giving more importance to the normative theory. This path can be compared to the maximum posterior trajectory (S13). While the linear interpolation follows a straight line (cyan), the MP estimates follow a curved trajectory (purple). While on many situations the two trajectories can be similar, sometimes there is a substantial difference in the trade-off, see Fig. S3A-C. In addition to the better trade-off, maximum entropy optimization priors also yield posterior distributions over parameters, Fig. S3D, which can be used for more refined analyses.

#### Interpolation using a Gaussian prior

Keeping the Bayesian approach, one could choose a family of optimization priors other than maximum entropy (MaxEnt). Here we consider Gaussian priors around a utility peak, with two diffferent levels of detail. The comparison with MaxEnt is in Fig. S4.

**Figure S3:**
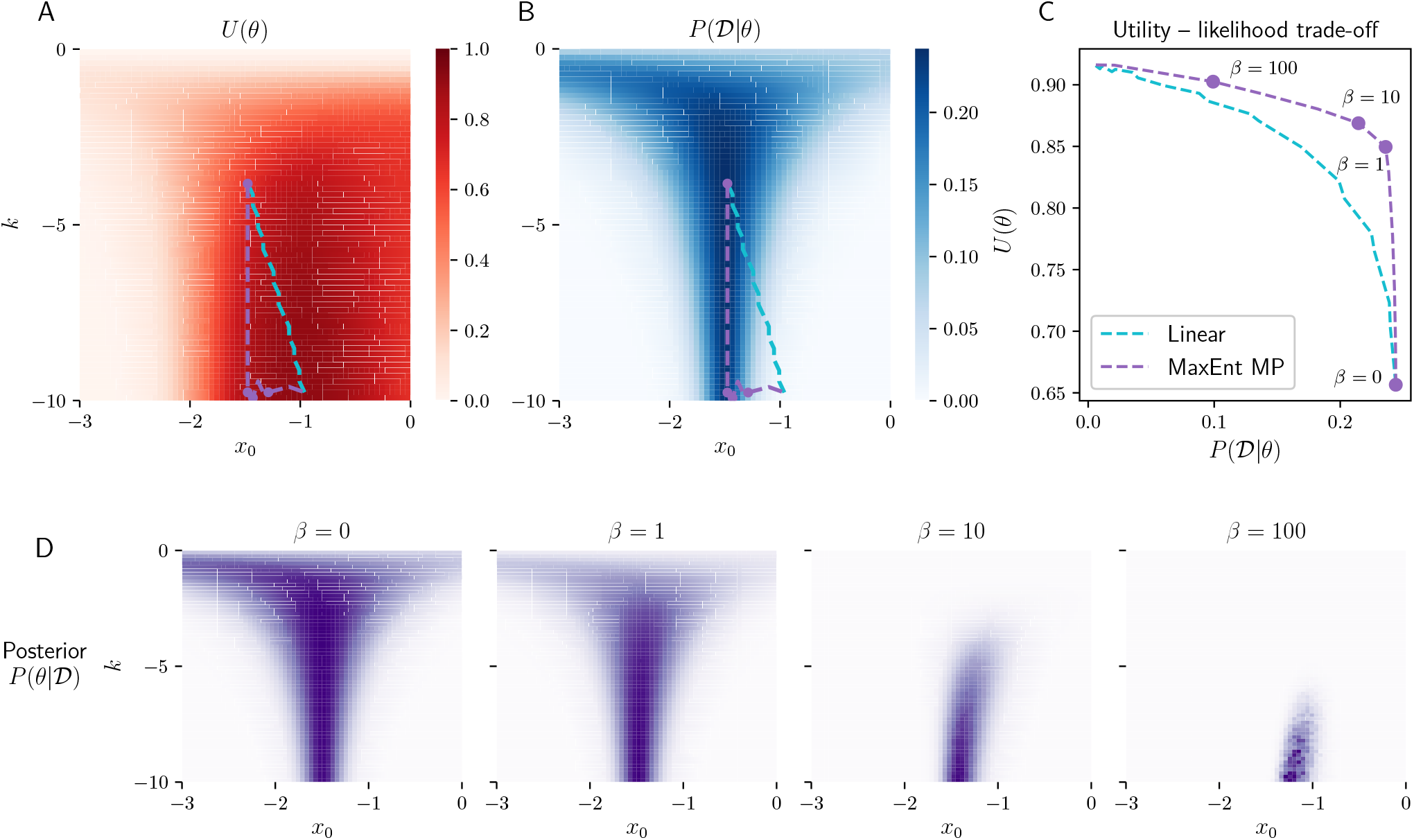
Interpolation between data and a normative theory in the parameter space. Linear interpolation takes a straight trajectory from maximum likelihood 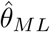 to maximum utility *θ*^∗^, cyan dashed lines. Maximum posterior (MP) interpolates between the same two, but takes a curved trajectory, purple dashed lines. **A-B**: Interpolation trajectories are shown on top of a utility heatmap (A) and a likelihood heatmap (B). **C**: MP achieves a better trade-off between likelihood and utility along the way. **D**: Posterior heatmaps, for the 4 values of *β* highlighted as purple points in the upper plots. Notes: The trajectories are bumpy, because parameters are rounded to the nearest vertex in a 128 by 128 grid, and the utility values are computed with noise (mutual information estimates from 200,000 samples). Data was chosen to get a marked difference between linear interpolation and maximum posterior trajectories. Likelihood is based on stimuli *x* = {*-*1.5, *-*1.49, 0, 2} and responses *r* = {0, 1, 0, 0}. For randomly generated data, the two trajectories are often similar in the utility-likelihood plot (even if they differ visibly in the *x*_0_, *k* plot).

The simplest option seems to be a Gaussian prior centered at the utility peak,

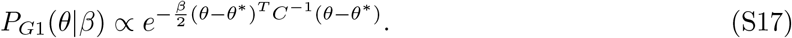

As an example using our toy model, we chose the covariance matrix *C* naively such that the Gaussian is roughly symmetric in the plots, corresponding to a visual “distance from the optimum”.

A more elaborate approach is to take into account the shape of the peak – the rate at which utility decreases in different directions. This is similar to the approach of Pérez-Escudero et al. [4]. We can fit the utility function around the peak with a quadratic function, and take the exponential to obtain a Gaussian,

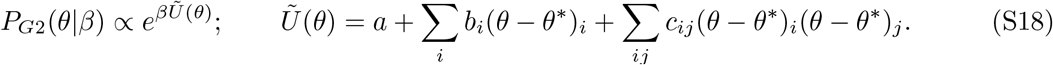

The linear coefficients *b*_*i*_ can be set to zero if the peak is inside the parameter domain – in our case the peak lies at the boundary, and utility has nonzero gradient there, and hence *b*_*i*_ will be nonzero. The coefficients *a, b, c* can be obtained by Taylor expansion if a formula for *U* (*θ*) is available; in our case *U* is computed numerically – so we fit *a, b, c* by minimizing mean square error in the vicinity of the peak.

Both types of Gaussian priors are parametrized by *β*, analogously to MaxEnt. Example priors for *β* = 10 are shown in Fig. S4A-C. To compare them in terms of the utility-likelihood trade-off, we compute the MP trajectory 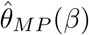 for increasing *β* as in the previous section. The trajectories are shown on top of each prior and on top of the likelihood surface, Fig. S4D. The panel S4E shows the utility-likelihood trade-off.

The trade-off is poor for the naive Gaussian *P*_*G*1_(*θ*|*β*). The fitted Gaussian prior *P*_*G*2_(*θ*|*β*) and the full MaxEnt prior achieve similar trade-offs for high *β* (i.e. near the peak), but *P*_*G*2_(*θ*|*β*) underperforms MaxEnt for lower *β*. As argued in the previous section, the trade-off is optimal for MaxEnt. However, the fitted Gaussian prior may often serve as a convenient approximation to reduce computational costs, especially if the utility has a unique maximum.

#### Discussion

This optimal trade-off achieved by the maximum entropy optimization priors is due to its sensitivity to the full shape of the likelihood and utility function – not only their peaks. In addition to the specific shape of the trajectory, this means that if the utility has multiple peaks, they are naturally taken into account.

The linear and Gaussian require specifying the “correct” peak, which can be difficult in high-dimensional systems. Consider the RF inference in Fig. 6C in the main text. Small values of *β* improve inference by pulling towards the nearest local maximum and gradually increasing utility. Too large *β* can lead the inference towards a different maximum with higher utility, which is however inconsistent with data. Linear interpolation towards this “wrong” maximum would yield meaningless results.

Except in situations when the data points to a point near a unique utility maximum, we need to take into account the full shape of the utility function (and of the likelihood function).

## S5 Data disambiguates degenerate theoretical predictions

Predictions of an optimization theory can be degenerate or ambiguous. Here we explore the *first kind of ambiguity*, where the utility function has multiple (possibly degenerate) maxima.

In this situation, biological context typically forces us to choose between two interpretations. On the one hand, we may observe multiple instances of the biological system and each instance could be an independent realization sampled from any of the maxima: statistical analyses of optimality thus need to consider and integrate over the whole parameter space, as in the approaches described above. On the other hand, we may observe a single (e.g., evolutionary) realization of the biological system which we hypothesize corresponds to a single optimum of the utility function. Our task is then first to identify that relevant maximum; if it exists, subsequent analyses can follow up on how well data agrees with that prediction and how surprising such an agreement might be in face of multiple alternative maxima.

**Figure S4:**
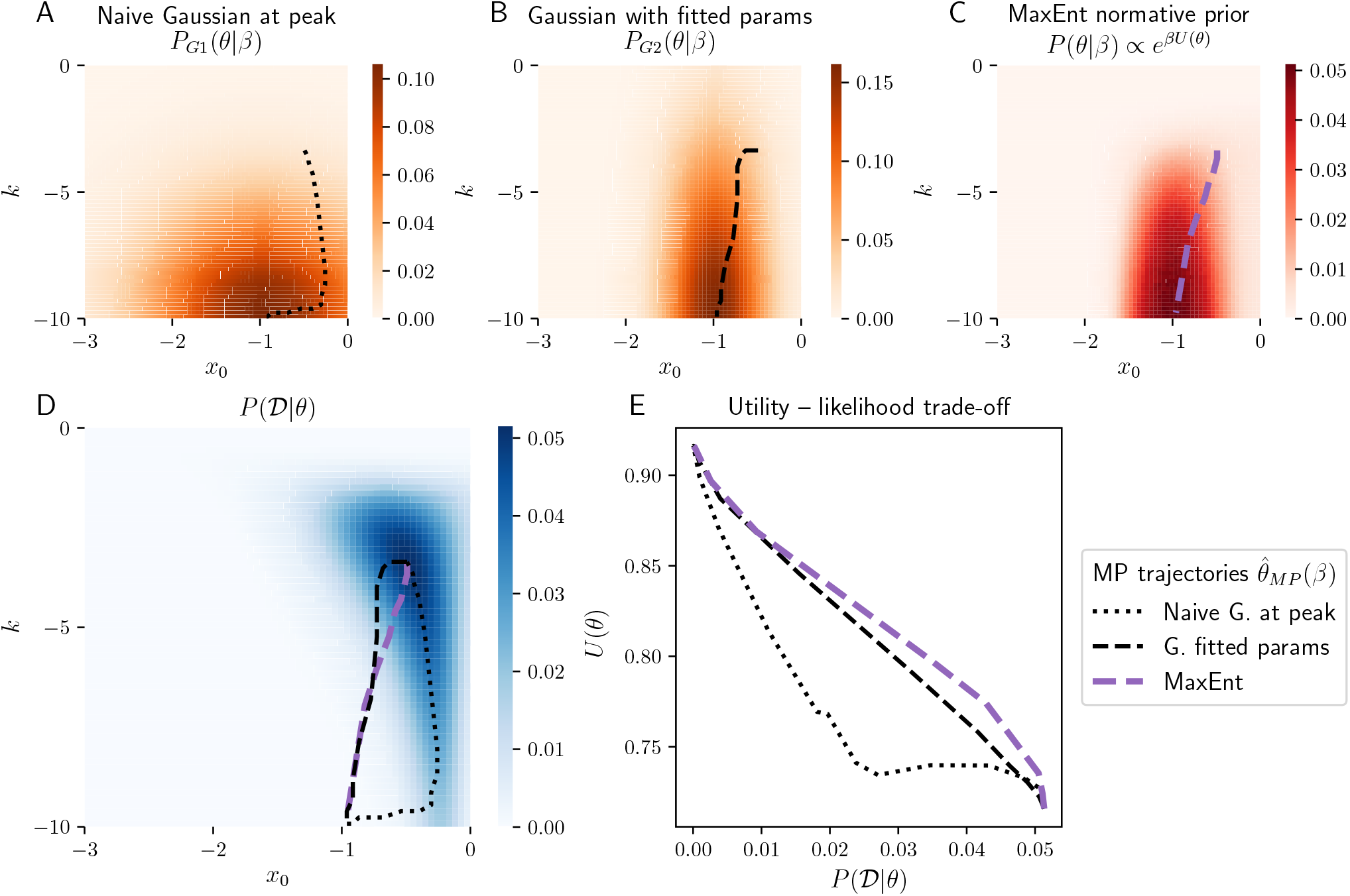
**A-C:** Alternative normative priors with *β* = 10. A: Naive Gaussian *P*_*G*1_(*θ*|*β*), see Eq. (S17). Covariance is *C* = ((10^2^, 0), (0, 3^2^)) such that the Gaussian is visually symmetric. B: Fitted Gaussian *P*_*G*2_(*θ*|*β*), see Eq. (S18). Parameters are *a* = 0.916, *b* = (*-*0.021, 0.212), *c* = ((0.003, *-*0.030), (*-*0.030, *-*0.603)) obtained from a least squares fit around the peak. C: MaxEnt prior *P* (*θ*|*β*) ∝ *e*^*βU*(*θ*)^. **D**: Likelihood surface used for further analysis. Generated from an intermediately optimised ground truth parameters and 20 spikes; same as in main text Fig. 3 likelihood 2. The trajectories (also shown in top panels) correspond to the maximum posterior 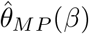 with varying *β* in the three priors above. **E**: Utility-likelihood trade off. MaxEnt performs best, and *P*_*G*2_(*θ*|*β*) is an approximation to it that performs similarly for high *β* (high utility region of the plot). The naive Gaussian *P*_*G*2_(*θ*|*β*) performs poorly, since it does not correspond to the shape of the utility peak.

In the Fig. 2 example of the main paper, multiple values of slope and offset yield optimal or close to optimal neural performance, resulting in ambiguous theoretical predictions. As a simple illustration of how data can break such ambiguities, we consider three example neurons with varying degree of optimality (Fig. S5A) and observe how their posteriors look like after seeing as few as *T* = 12 stimulus-response pairs from each neuron (Fig. S5B). All three simulated datasets reduced the uncertainty (entropy) about the neuron’s parameters by a similar amount, as reflected by the entropy and utility of the posterior versus the entropy and utility of the prior (Fig. S5C). Despite similar reductions in entropy, the resulting inferences were very different in terms of agreement with the theory. Only the posterior of the first neuron concentrated in a high-utility region of the parameter domain, thus clearly identifying one of the four peaks of the utility function as consistent with the operating regime of the simulated neuron. The two remaining posteriors are concentrated in regions of the parameter space which weakly overlap with the prior, or where prior probability is close to 0. To capture these qualitative differences mathematically, we define and compute the *mode entropy*, where each mode corresponds to the attraction basin of a local utility maximum. Optimality theories with degenerate maxima will allocate the prior probability relatively evenly among the modes, resulting in high mode entropy (here, 2 bits, i.e., 4 possible local maxima). A few observations of neuron 1 consistent with an optimal solution drastically collapsed this mode uncertainty and identified the single relevant utility maximum; this decrease was smaller for slightly suboptimal neuron 2 and vanished for neuron 3 (Fig. S5D).

This is a very non-standard application of the Bayesian framework at small sample sizes, *T* : here, the structure of the prior (i.e., the normative theory) dominates the posterior, in what we refer to as the “data-regularized prediction” regime. In this regime, our goal is to derive *ab initio* theoretical predictions, not fit parameters to reproduce the data, and the data is only used to disambiguate the prediction – to identify which utility maximum, if any, is realized in nature. If we track the evolution of the average utility, full posterior entropy, and the mode entropy with the number of data points *T*, we clearly see the transition from such “data-regularized prediction” regime dominated by the prior normative theory, to the “theory-regularized inference” regime in the large sample limit (Fig. S5E). In the first regime, data removes the theoretical ambiguity and collapses the mode entropy with *T <* 10 samples; in the second regime, the actual parameter values (*k, x*_0_) are inferred with increasing precision, as evidenced by posterior entropy that continues to decrease linearly in the log sample size (corresponding to the standard asymptotic inverse scaling of the variance in parameter estimates with the sample size).

In the “data-regularized prediction” regime, *β* also serves a novel role: when the normative theory has multiple optima with a broader spectrum of utility values, *β* determines which of the peaks are considered as nearly degenerate candidate predictions. A peak with utility *U* ^*0*^ *< U*_max_ will be suppressed in the prior by ∼exp(- *β*(*U*_max_ *U*′)), and, for sufficiently high *β*, the alternative theoretical prediction corresponding to *U* ^*0*^ will be disregarded irrespective of the data.

Here we showed that the ambiguities of normative theories resulting from degenerate utility maxima can often be resolved in our Bayesian framework in the “data-regularized prediction” regime by a very small amount of data. This power may appear trivial at first glance, because the parameter space of our example is two dimensional and so priors and posteriors can be evaluated explicitly and plotted across their whole domain. In more realistic cases involving tens of parameters, however, finding all (nearly) degenerate maxima of the utility function and deciding whether data is “close to” any one of them becomes a daunting task due to the curse of dimensionality. In the past, this has severely limited the application of optimality principles to complex systems with more than a few parameters [5]–[7], except in those rare cases where strict guarantees exist [8]. In contrast, even in spaces of high dimensionality, posteriors resulting from our framework can be sampled with Monte-Carlo methods or optimized by well-developed methodology [9], with search concentrated around the unique peak of the normative theory that is simultaneously permitted by the chosen value of *β* and is consistent with the data, if such a peak exists. Intuitively, theory “proposes” possible optimal solutions *ab initio* while data “disposes” with those degenerate solutions for which there is no likelihood support.

## S6 Sparse and slow utility in V1 receptive fields

To complement the analysis of V1 neurons in terms of the sparse utility *U*_SC_, here we present additional analysis in terms of the slowness utility. Slowness utility *U*_LC_ assumes that neurons extract invariant properties of sensory data [10]. Given a linear filter *c*, we quantified slowness of its responses to a set of natural image sequences using the following function:

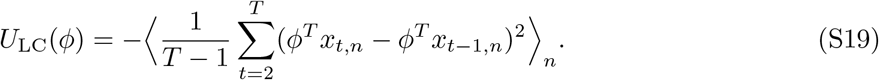

where *n* is an index over image sequences, and *t* is a time index over images within a sequence. Filter slowness was averaged across a 5 *·* 10^4^ artificially generated natural image sequences of length *T* = 2. Each sequence was generated by moving an image patch by a random distance *n*_*x*_ ∈ [*-*8, 8] pixels in a horizontal direction and *n*_*y*_ ∈ [8, 8] pixels in vertical direction, and rotating it by a random angle *α* ∈ [90^*0*^, 90^*0*^]. The mean and standard deviation of filters *c* and image patches *x*_*t,n*_ was set to be 0 and 1 respectively.

Optimally slow RFs minimize temporal variability of neural activity in natural sensory enviromnents [11]. On the level of individual neurons, slowness and sparseness optimality criteria yield very different predictions. In contrast to optimally sparse RFs which are localized in space and frequency (Fig. S6B, left column), RFs optimized for slowness are broad and non-local (Fig. S6B, right column).

In the right column of Fig. S6B-F, we present analysis of the optimality of V1 RFs in terms of the slow utility *U*_LC_. This analysis is complementary to the analysis of the sparse utility *U* presented in the main text. We present analysis of the sparse utility again in the left panels of Fig. S6B-F for comparison. To compute posteriors over *β* (Fig. S6D), for slow utility we used 64 *β* values equally spaced on the [-32, 32] interval. The remaining details of the analysis are shared with the analysis of sparse coding utility and are described in the Methods.

Overall, RFs of individual neurons are much more consistent with the sparse, rather than with the slow utility. This is readily apparent in the outcome of the optimality test (Fig. S6 C) and posteriors over the utility parameter *β* (Fig. S6 D), where V1 RFs yield a slightly negative estimate 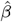. Similarly, the empirical distribution of utility values of V1 RFs as well as their spatial autocorrleation, is more consistent with the inferred distribution of sparse, rather than slow utilities (Fig. S6E and F respectively).

Here we analyzed utility of *indiviudal* neurons, treating them as independent realizations from an underlying distribution of parameters. It is important to stress that simultaneous optimization of a *population* of model neurons for maximal slowness yields filters which very closely resemble RFs of visual neurons [11], [12]. Moreover, slowness and sparseness are both affected by eye movements and natural stimulus dynamics, while the RFs used here were recorded in anesthesized and paralyzed animals. Our analysis is therefore not a proof of lack of optimization for slowness at the population level. It is rather a demonstration of applicability of the framework to real data. Analysis of optimality of neural populations is a subject of future work.

**Figure S5:**
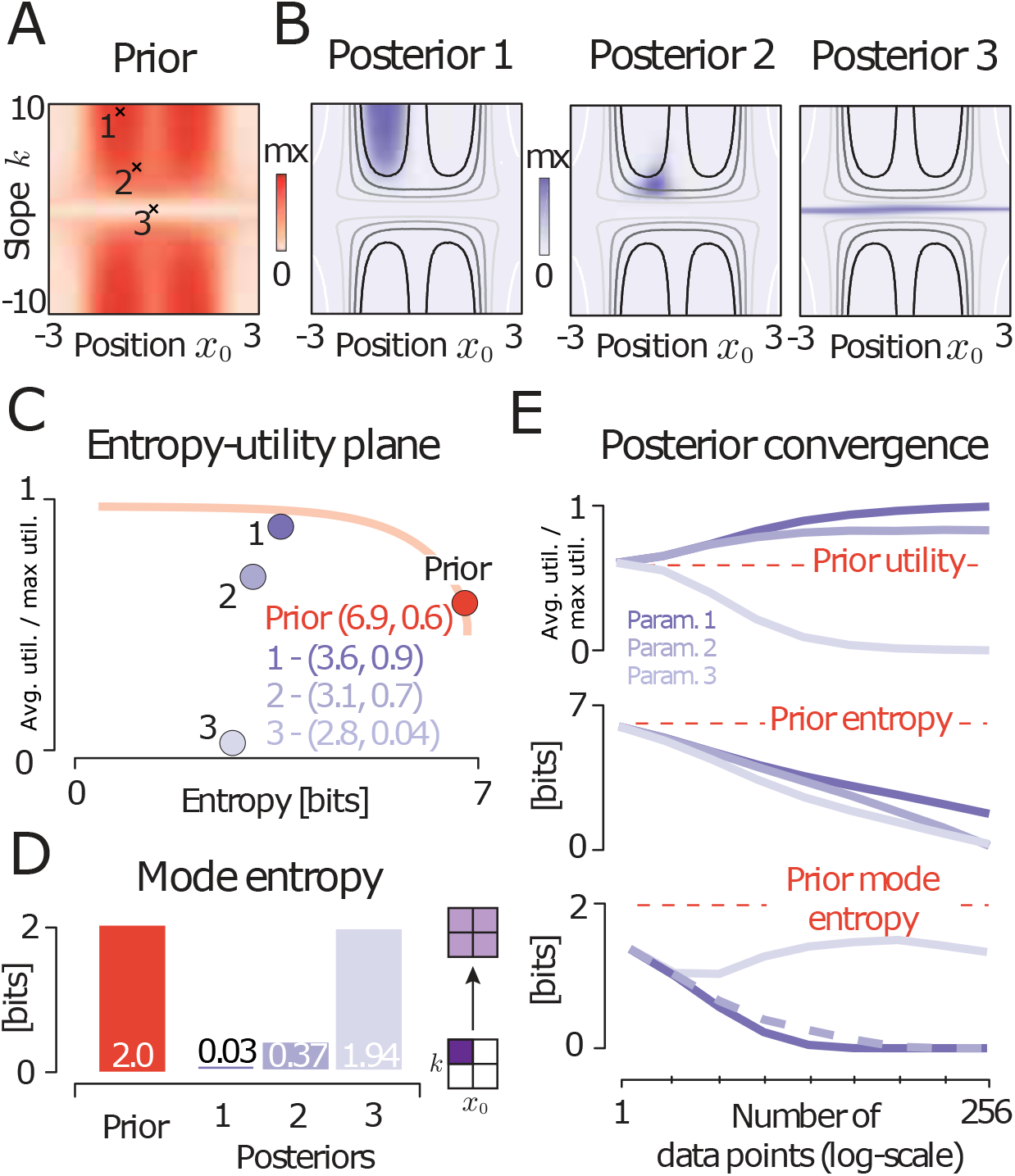
Disambiguating degenerate theoretical predictions. **(A)** A maximum-entropy prior derived from the mutual information utility with *β* = 1. The prior has multiple maxima reflecting non-uniqueness of theoretical predictions. **(B)** Posteriors obtained by updating the prior with three example datasets (𝒟_1_, 𝒟_2_, 𝒟_3_). Grayscale lines denote regions of different utility values (black – highest utility, white – lowest utility). Depending on the observed data, posteriors concentrate in regions of different utility value. **(C)** Distributions on the entropy-utility plane. Orange dot corresponds to the prior from A, purple dots to posteriors from B. Orange line is the entropy–average utility tradeoff in the maximum entropy optimization prior (analogous to Fig. 2E in the main text). **(D)** Mode entropy. In the prior (red bar), probability is equally distributed across 4 peaks of the distribution resulting in 2 bits of entropy. Mode entropy decreases significantly in posteriors 1 and 2. **(E)** Posterior convergence. Average utility (top row), posterior entropy (middle row) and posterior mode entropy (bottom row) are plotted against the number of data samples; shown are averages of 512 realizations for each data set size. Purple lines correspond to parameter settings 1-3 in panel A. Red dashed line denotes values of each statistic for the prior.

**Figure S6:**
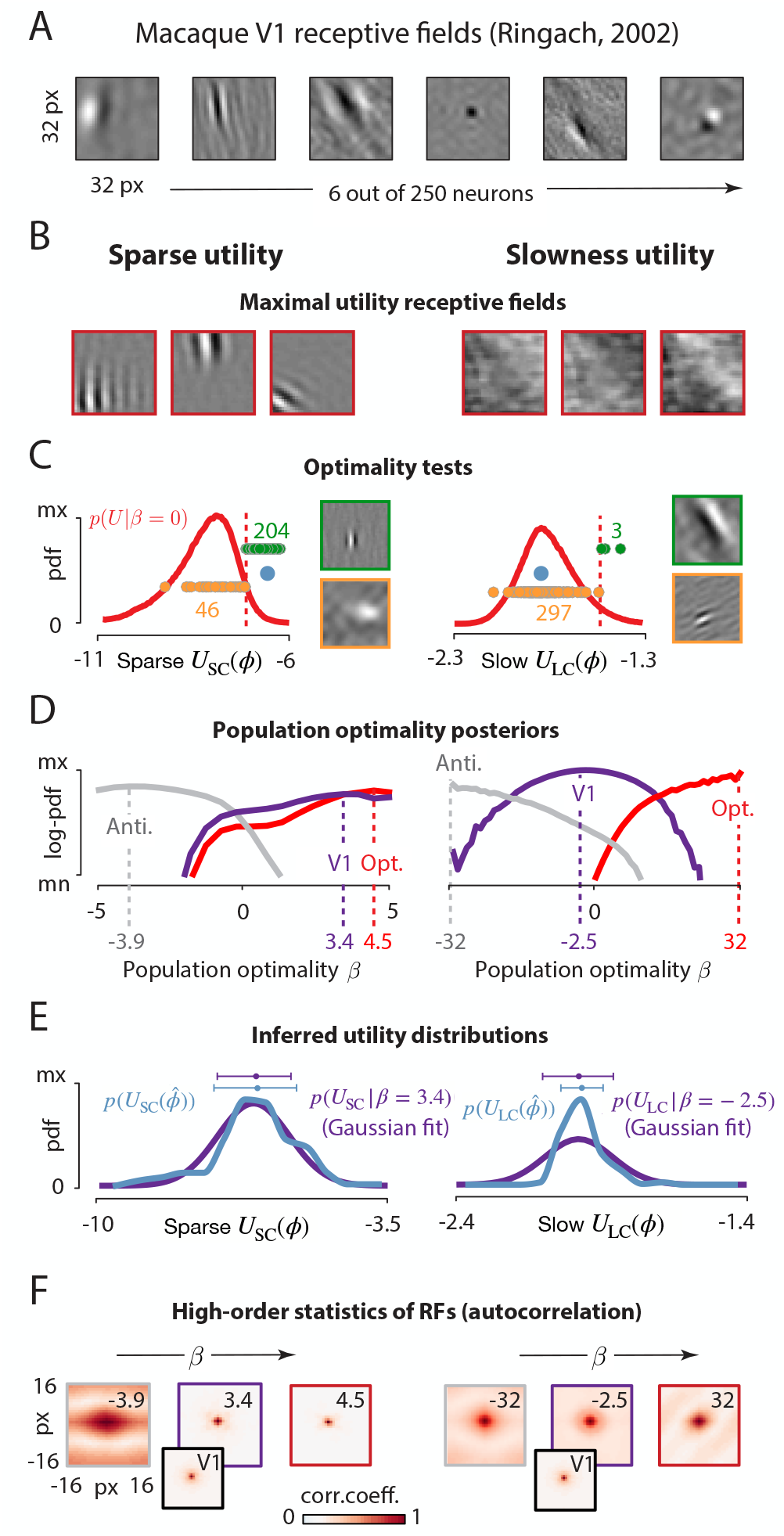
Sparse and slow utility analysis of V1 receptive fields. **(A)** Six example receptive fields (RFs) from Macaque visual cortex (courtesy of Dario Ringach; [13]). **(B)** Simulated RFs optimized for sparsity (left column) and slowness (right column). **(C)** Null distributions of utility values used to test for optimality under sparse (left column) and slow (right column) utilities. Red dashed lines denote the significance threshold (95th percentile). Green and orange circles correspond to significant and non-significant receptive fields (the axis was truncated for visualization purposes, and not all values are displayed). Example significant and non-significant receptive fields are displayed in green and orange frames respectively. Blue dots show the average utility of receptive fields, which are equal to the 99.6^th^ percentile (sparse *U*_SC_) and 46^th^ percentile (slow *U*_LC_) of *p*(*U*|*β* = 0). **(D)** Approximate log-posteriors over population optimality parameter *β* derived from 250 RFs estimates (purple line), 250 maximum-utility filters (red line) and 250 minimal-utility filters (gray line). Dashed lines mark MAP estimates of beta. **(E)** Empirical distributions of RF utilities (blue lines) compared with utility distributions consistent with the population optimality *β* inferred from V1 data (purple lines). **(F)** Spatial autocorrelation of RFs consistent with different average values of utility (determined by *β* parameter). Values of *β* are denoted in the top-right corner of each panel, and correspond to results of inference displayed in panel D. Middle plots (purple frame) in the left and the right column depict autocorrelation consistent with *β* inferred from V1 RFs. For comparison, autocorrelation of RFs is displayed as an inset.

## Footnotes

1 We could also include a non-uniform null distribution *q*(*θ*) as discussed in Sec. S2, and the MP formula would be 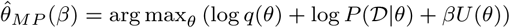

